# The role of UV-B light on small RNA activity during grapevine berry development

**DOI:** 10.1101/375998

**Authors:** Sukumaran Sunitha, Rodrigo Loyola, José Antonio Alcalde, Patricio Arce-Johnson, José Tomás Matus, Christopher D. Rock

## Abstract

UV-B regulation of anthocyanin biosynthesis in vegetative and grapevine berry tissues has been extensively described. However, its relation with UV-B-regulated microRNAs (miRNAs) has not been addressed before in this species. We explored by deep sequencing of small RNA libraries the developmental dynamics and UV-B effects on miRNAs and associated phased small interfering RNA (phasi-RNAs)-producing loci abundances in *in vitro*-grown plantlets, in field-grown berry skins of cv. Cabernet Sauvignon, and low- and high UV-B fluence treatments of greenhouse-grown berries at several time points around veraison. We observed by RNA blotting a differential effect of low-versus high-fluence UV-B on miR828 abundances (an effector of anthocyanins and UV-absorbing polyphenolics) across berry development, and identified other miRNAs that correlated with miR828 dynamics. The functional significance of the observed UV-coordinated miRNA responses to UV was supported by degradome evidences of AGO-programmed slicing of mRNAs. Inverse co-expression of the up-regulated miRNAs miR156, miR482, miR530, and miR828 with cognate target gene expressions in response to high fluence UV-B measured by quantitative real-time PCR. These UV-response relationships were also corroborated by analyzing three published transcriptome datasets (berries subjected to UV-C for 1 hr [at pre-veraison], UV-B for five weeks post-veraison, and five red-skinned varieties across four berry development time points). Based on observed significant changes by UV-B on miRNA and derivative phasi-RNA abundances, we propose a regulatory network model of UV responses impacting anti-oxidant and stress-associated polyphenolic compound biosynthesis. In this model high-fluence UV-B increases miR168 (validated in a UV-B small RNA-derived degradome library to target *ARGONAUTE1*, which spawns phasi-RNAs) and miR530 (targets a novel Plus-3 domain mRNA), while decreasing miR403 abundances (validated to target *ARGONAUTE2*), thereby coordinating post-transcriptional gene silencing activities by different AGOs. Up-regulation of miR3627/4376 (validated to target Ca^2+^-transporting ATPase10 that spawns phasi-RNAs) could facilitate anthocyanin accumulation. miR395 and miR399, induced by sulfur and phosphorus starvation in other species (conditions known to trigger anthocyanin accumulation) respond positively to UV-B radiation and are shown to slice cognate targets in grapevine. miR156/miR535 is shown to target *SQUAMOSA PROMOTER-BINDING* transcription factor genes that potentially regulate the activities of MYB-bHLH-WD40 complexes and thereby anthocyanin biosynthesis. Increases in MYB-bHLH-WD40 TFs could also contribute to the observed up-regulation of miR828 via the conserved and degradome-validated auto-regulatory loop involving miR828/*TAS4abc* to regulate *MYBA6/A7/A5-MYB113-like* levels and thereby anthocyanin levels. These results and meta-analysis provide a basis for systems approaches to better understand non-coding RNA functions in response to UV.

## 1 Introduction

The sessile nature of plants makes them vulnerable to myriad abiotic and biotic stress factors to which they must adapt to survive. Ultraviolet (UV) irradiation, drought, heat, cold, salt, and oxygen deficiency are the major abiotic factors restricting plant growth, development, and productivity whereas microbial pathogens and herbivores are major biotic stress factors. Genome-wide association studies have identified UV-B responses with autoimmune-like reactions due to overexpression of defense related genes (Piofczyk, Jeena, & Pecinka, 2015). UV-B radiation (280-315 nm) is mostly absorbed by the ozone layer although about two percent passes through the atmosphere and causes detrimental effects on all life forms. A highly-conserved UV-B perception and signaling system is evolved in plants (Heijde & Ulm, 2012; Tilbrook *et al*., 2016). Exposure to low fluence doses of UV-B irradiation acts as an environmental trigger facilitating expression of wide array of genes involved in development such as auxin, ethylene, abscisic acid (ABA), and cell wall modification (Frohnmeyer & Staiger, 2003; Jenkins, 2017; Mackerness, 2000; Pontin *et al*., 2010; Ries *et al*., 2000). In addition to the up-regulation of antioxidant/free radical scavenging enzymes and proteins involved in DNA repair and cell cycle proteins by UV (Brosché, Schuler, Kalbina, Connor, & Strid, 2002), a highly specific signaling pathway involving photomorphogenic receptors, some of which influence miRNA activities, is well characterized (Jenkins, 2017). In Arabidopsis, Cryptochrome1 and Cryptochrome2 mediate the expression of miR172 after blue light stimulation in a CONSTANS-independent manner to regulate photoperiodic flowering time (Jung *et al*., 2007). miR408 is coordinately regulated by SQUAMOSA PROMOTER BINDING PROTEIN-LIKE7 (SPL7) and ELONGATED HYPOCOTYL5 (HY5) in response to light (Zhang *et al*., 2014).

Plants fend off harmful effects of UV-B with plant-specific phenolic secondary metabolites derived from phenylalanine and deposited in the vacuoles of epidermal cells (Dotto & Casati, 2017; Li, Ou-Lee, Raba, Amundson, & Last, 1993). Flavonols, chalcones, and anthocyanins are a diverse group of polyphenolic secondary metabolites synthesized by coordinated transcriptional control of a suite of biosynthetic enzymes including *CHALCONE SYNTHASE* (*CHS*). UV-B induction of *CHS* (Jenkins, 2017) and the downstream phenylpropanoid biosynthetic genes is well characterized. UV RESISTANCE LOCUS 8 (UVR8), a UV-B receptor, upon absorbing UV-B radiation monomerizes into active form (Rizzini *et al*., 2011; Wu *et al*., 2012) and interacts with CONSTITUTIVELY PHOTOMORPHOGENIC1 (COP1) (Cloix *et al*., 2012). COP1 is a WD40/RING protein that facilitates proteasome-mediated degradation in the dark of HY5, a basic leucine zipper transcription factor (TF). Thus, COP1 functions as a central mediator of UV-B adaptation and light photomorphogenic responses (Saijo *et al*., 2003). COP1 interaction with UVR8 releases HY5 from proteasome-mediated degradation and promotes photomorphogenic signaling (Huang *et al*., 2013). HY5 binds to *CHS* promoter, promoter elements of MYeloBlastosis (MYB) transcription factor (TF) *MYB12*, a regulator of flavonol biosynthesis (Lee *et al*., 2007, Stracke *et al*., 2010), and *MYB75/PRODUCTION OF ANTHOCYANIN PIGMENT1 (PAP1)*, a TF inducer of anthocyanin biosynthesis genes (Shin *et al*., 2013).

Despite grapevine (*Vitis vinifera* L.) having a sequenced genome available for over a decade and being an important fruit crop abundant in health-promoting- and flavor-enhancing polyphenolics, UV-B signaling components and their functions in integrated control of secondary metabolism were not identified until recently. Loyola *et al*. (2016) characterized the grapevine photomorphogenic factors *UV-B RECEPTOR* (*UVR1*), *HY5* and *HY5 HOMOLOGUE* (*HYH*) that function in UV-B perception and signaling. The expression profile of the above-mentioned photomorphogenic factors was studied in vegetative and reproductive tissues at different developmental stages. While *UVR1* is strongly expressed in early berry developmental stages, *UVR1* transcript abundance is mostly influenced by light and temperature but not UV-B radiation. *HY5* is induced by low fluence UV-B in leaves and at the early berry developmental stages (both low and high fluence) (Loyola *et al*., 2016). *HYH* is induced by high fluence UV-B three weeks after the onset of ripening, (veraison, manifested as color change of berries and a transition from organic acid to sugar accumulation). UV-B irradiation also promotes flavonols accumulation in leaf and berries. Thus, both high and low UV-B exposures in grapevine may facilitate flavonol accumulation in vegetative and reproductive organs through the activation of *HY5* and *HYH* TFs at different developmental stages (Matus, 2016).

The grapevine berry color locus in chromosome 2 encodes several R2R3-type MYB genes including the anthocyanin regulators *MYBA1* and *MYBA2* (Walker *et al*., 2007; Rinaldo *et al*., 2015). The red pigmentation observed in vegetative tissues is attributed to activity of *MYBA6, MYBA7*, and potentially *MYBA5* genes on chromosome 14 (Matus *et al*., 2017). *MYBA5/6/7* accumulate at high levels in early development of berries and decrease after the onset of veraison. UV-B radiation induces *MYBA1* genes and significantly delays the down-regulation of *MYBA6* and *MYBA7* at the latter stages of berry development (Matus *et al*., 2017). Interestingly, *MYBA6* and *MYBA7* follow the exact expression pattern of grapevine *HY5* (Loyola *et al*., 2016), suggesting *MYBA6* and *MYBA7* also contribute to fruit pigmentation upon increased radiation stress, in addition to *MYBA1*.

Grapevine *MYBA6* and *MYBA7* are orthologs of *Arabidopsis* MYB TFs *PAP1/MYB75* and *PAP2/MYB90* (Borevitz, Xia, Blount, Dixon, & Lamb, 2000). *PAP1/MYB75, PAP2/MYB90* and *MYB113* are under negative regulatory control of *Trans Acting Small-interfering RNA locus4 (TAS4)* that generates a 21 nt antisense trans-acting small-interfering RNA (tasi-RNA) species TAS4-3′D4(-) by the processive endonuclease activity of DICER1/4 (Rajagopalan, Vaucheret, Trejo, & Bartel, 2006). *TAS4* siRNA production requires miR828 programming of ARGONAUTE (AGO)-containing RNA Induced Silencing Complex (RISC). miR828 loaded in RISC binds to its complementary site in *TAS4*, directs cleavage and sets the phase for 21-nt siRNA production in association with RNA-DEPENDENT RNA POLYMERASE 6. The fourth antisense phase tasi-RNA species generated by miR828-mediated cleavage is *TAS4*-3′D4(-) siRNA. A RISC programmed with *TAS4*-3′D4(-) siRNA binds to complementary sequences in *AtPAP1/PAP2/MYB113* mRNAs, which undergo endonucleolytic slicing (Rajagopalan *et al*., 2006). In addition to *TAS4*-3′D4(-) predicted cleavage, *AtMYB113* is also targeted by miR828 directly, suggesting a close evolutionary relationship between *AtPAP1/PAP2/MYB113*, miR828, and *TAS4* (Rajagopalan *et al*., 2006). *TAS4* is functionally conserved among various eudicots (Luo, Mittal, Jia, & Rock, 2012; Rock, 2013) and TAS4-3′D4(-) targets grapevine *VviMYBA6* and *VviMYBA7* which in turn triggers 21 nt phased small interfering RNA (phasi-RNAs) production from *VviMYBA6* and *VviMYBA7* in register with the TAS4-3′D4(-) slicing site (Rock, 2013; Zhang, Li, Wang, & Fang, 2012). The biological significance of 21 nt phasi-RNAs is largely unknown, but hypothesized to be important for defining a concentration gradient for silencing activity across cell layers and for widespread *trans* suppression or homeostasis of highly conserved motifs in the large families of Leucine Rich Repeat (LRR) resistance genes (Fei, Xia, & Meyers, 2013) in response to pathogen effectors (Shivaprasad *et al*., 2012), or for diversification and neofunctionalization of the large Pentatrico Peptide Repeat (PPR) family of genes (Howell *et al*., 2007).

Studies to date have focused on the influence of UV-B radiation in regulating anthocyanin biosynthesis at different developmental stages of berry development. The effect of UV-B on miRNA profiles is well-documented in *Arabidopsis*, poplar, maize and wheat (Zhou, Wang, & Zhang, 2007; Jia, Ren, Chen, Li, & Tang, 2009; Casati, 2013; Wang *et al*., 2013), however the role of UV-B in post-transcriptional regulation of *MYB* TFs by miR828/TAS4 pathway is unexplored. Further, there is a large annotation gap between the empirical knowledge of siRNA expression, especially the role of miRNAs and tasi-RNAs on production and functions of phasi-RNAs (Coruh, Shahid, & Axtell, 2014; Luo *et al*., 2009). This work focuses on a data-driven systems approach to explore UV-B effects on miRNA/phasi-RNA expression and elucidate functional consequences for stress adaptation by controlling the biosynthesis of UV-B absorbing anti-oxidant secondary metabolites. We performed a detailed computational re-analysis of empirically characterized grapevine miRNAs for slicing activities with a degradome library (Pantaleo et al., 2010). We then performed quantitative real-time PCR (qRT-PCR) assays on miRNA target mRNAs in same-source samples used to generate our small RNA (sRNA) libraries and obtained supporting evidence for a regulatory network model of UV responses mediated by sRNAs, further substantiated by meta-analysis of available mRNA transcriptome results for similar experiments across UV treatments and developmental time points (Carbonell-Bejerano *et al*., 2014; Massonnet *et al*, 2017; Suzuki *et al*., 2015).

## 2 Materials and Methods

### 2.1 UV-B treated *in vitro* plantlets and fruits

In order to gain perspective in defining berry-specific relationships (Loyola *et al*., 2016; Czemmel *et al*., 2017) relative to leaf UV-B (Cavallini *et al*., 2015) sRNA responses, Cabernet Sauvignon cultivar (‘cv’) *in vitro* grown plantlets (four negative control and four UV-B treated biological replicates; n=4) were exposed to 6 hr of low fluence UV-B radiation (~0.15 W m^-2^ irradiance) exactly as conducted in Cavallini *et al*. (2015). Leaves were harvested after +UV-B treatment and frozen in liquid nitrogen for RNA extraction. Control plants were grown under the same conditions described above but covered with a polyester filter (100-μm clear safety polyester plastic film) that absorbed total UV-B from the spectrum without affecting the photosynthetic active radiation.

Potted nine-year-old greenhouse-grown cv. ‘Cabernet Sauvignon’ fruit clusters were exposed to high (~ 0.3 W m^-2^ daily for 5 hr) UV-B or low (~ 0.1 W m^-2^ daily for 10 hr) UV-B irradiance during 2011-2012 and 2012-2013 growing season, respectively (both contrasted to control ‘filtered UV-B’ conditions; more details in Loyola *et al*. 2016). Sample collection was carried out at four stages of grapevine development: −3, 0, 3 and 6 weeks before (-) and after (+) veraison (WAV, defined as the time at which clusters are 30-50% colored and sugar concentration reaches 5° Brix). Berries were immediately peeled, deseeded and skins were frozen in liquid nitrogen for RNA extraction. A total of 18 berries were sampled every 3 weeks from 9 grape clusters (n=3, three bunches per plant and two berries per cluster). One and three biological replicates were considered for deep sequencing and qRT-PCR analyses, respectively.

A solar UV-B filtering treatment was applied in a commercial cv. ‘Cabernet Sauvignon’ field, located in the Maipo Valley (Chile; 33° 43′ 28″ S, 70° 45′ 9.71″ W) during 2011-2012 growing season exactly as conducted in Czemmel *et al*. (2017). Berry skin samples were collected at −3, 0, +3 and +6 WAV from a row of UV-B filtered (UV-B radiation was blocked by installing a 100-μm clear polyester film at the position of grape clusters) and +UV-B irradiated plants (natural solar radiation, no filter). The experiment was designed in four blocks (rows) with five plants each (biological replicates *n* = 4). Three berries per cluster (randomly sampled) and four clusters per plant were used for each sample. One biological replicate was considered for deep sequencing.

In total, 28 samples were used for sRNA library construction; four for *in vitro* plantlets (two biological replicates, two conditions), 16 for greenhouse-grown berries (one biological replicate, four time points, four conditions [with and without high- or low-UV-B fluence), and eight for field-grown berries (one biological replicate, four time points, two conditions [with or without solar UV-B]).

### 2.2 RNA extraction and sRNA libraries construction

Total RNA was isolated following the procedure of Reid, Olsson, Schlosser, Peng, and Lund (2006), using a CTAB-spermidine extraction buffer. The RNA was quantified using NanoDrop 1000 spectrometer (Thermo Fisher Scientific). The integrity of RNA was assessed by an Agilent 2100 Bioanalyzer using a sRNA chip (Agilent Technologies) according to the manufacturer’s instructions.

sRNA libraries were prepared using total RNA as input (1 μg) according to the instructions provided by TruSeq sRNA Sample Preparation Kit (Illumina®). Twenty-eight bar-coded sRNA libraries were constructed. The quality of each library was assessed using an Agilent High Sensitivity DNA chip on an Agilent 2100 Bioanalyzer. Equi-molar concentrations of libraries were pooled and sequenced on an Illumina NextSeq500v2 instrument at the Institute for Integrative Genome Biology, UC Riverside. One library (minus UV-B, −3 WAV, high fluence UV greenhouse experiment) was resequenced to increase reads depth on a MiSeqv3 platform, and another library was solely sequenced on MiSeqv3 (plus UV-B, +6 WAV, field grown; Supplementary Table 1a).

### 2.3 Bioinformatics and Statistical Analyses of Sequencing Data

The quality assessment of the raw reads was done by FastQCv0.11.5 (https://www.bioinformatics.babraham.ac.uk/projects/fastqc/) and adaptor sequences were trimmed using fastx_clipper (http://hannonlab.cshl.edu/fastx_toolkit/index.html). Reads longer than 18 nt were retained and were sequentially mapped to *Arabidopsis thaliana* rRNAs, tRNAs, snRNAs, and transposable elements (TEs; downloaded from https://plants.ensembl.org/info/website/ftp/index.html and www.arabidopsis.org) using Bowtie (Langmead, Trapnell, Pop, & Salzberg, 2009) to remove reads matching the same. The pre-filtered reads were mapped to the reference *Vitis vinifera* 12X genome sequence, version NCBI RefSeq GCF_000003745.3 (https://www.ncbi.nlm.nih.gov/assembly/GCF_000003745.3) (Jaillon *et al*. 2007) using ShortStack (version 3.8.2) (Johnson et al., 2016). In order to increase the power of ShortStack to call *MIRNAs* by sequencing at least one rare miRNA star species, we included publicly available sRNA datasets for grapevine from the following sources available from NCBI: Belli Kullan *et al*. (2015; SRR1528350-SRR1528419), Paim-Pinto *et al*. (2016; SRR4031591-SRR4031638), Pantaleo *et al*. (2010; 2016; GSM458927-GSM458930; GSM1544381-GSM1544384), Sun *et al*. (2015; SRR2029783, SRR2029784), Wang *et al*. (2012, 2014; SRR390296, GSM1907875-GSM1907880), and Zhai *et al*. (2011; GSM803800-GSM803803). The ShortStack counts output for these libraries were not used for subsequent differential expression analysis. ShortStack counts output generated from our UV treatment libraries alone were subjected to differential expression analysis.

A few novel candidate *MIRNAs* were called de novo by ShortStack as having met all annotation requirements (viz., > 50% of reads map to hairpin duplex, >80% of reads map to one strand of the hairpin, and at least one miRNA star species sequenced). The raw counts output of ShortStack for 40,678 sRNA-producing loci with greater than 261 total reads per cluster summed across all libraries was used as input to DESeq2 “R” Package (version 1.16.1) (Love, Huber, & Anders, 2014) to analyze the differential expression of sRNA cluster loci under different conditions. The cutoff was chosen to capture the low-expressed (~ 1.5 reads per million, rpm) *MIR828* and *MYBA6* and *MYBA7* clusters, which were of particular interest but at the cost of more independent statistical tests. For comprehensive genome-wide assessment, an additional 25 clusters mapping to miRBase21-annotated grapevine *MIRNAs* (Kozomara & Griffiths-Jones, 2014) were included which were below the cutoff of 261 total reads. This preprocessing step of limiting candidate clusters increases the power of DESeq by removing poorly expressed loci that would otherwise contribute to ‘shot noise’ variation and excessive multiple test corrections for statistical significance, while adequately encompassing nearly all annotated *MIRNA* loci (e.g. *MIR858*, 0.4 rpm; *MIR395f*, 0.05 rpm (see Supplementary Table 3).

The experimental comparisons made for differential expression analyses of biologically coordinated replicates are outlined in Table 1. The differential expression of *MIRNA* loci and phasi-RNA-producing loci as a time-series spanning berry development was tested using the likelihood ratio test (LRT) for inference, which assumes that the expression data follows negative binomial general linear model. To detect differentially expressed genes under +UV/-UV and high/low fluence UV radiation, Wald-Log test was performed. Publicly available grape degradome data (Pantaleo *et al*., 2010; GSM458931) and the sRNA library datasets were subjected to PhaseTank analysis (Guo, Qu, & Jin, 2015) to identify the genome-wide phased siRNA-producing loci and regulatory cascades triggered by miRNAs. PhaseTank is built on CleaveLandv4.4.3 (Addo-Quaye, Miller, & Axtell, 2009). Cleaveland4 is more sensitive than earlier versions employed in prior analyses of GSM458931 (Pantaleo *et al*., 2010) by implementation of Generic Small RNA-Transcriptome Aligner (GSTAr) which calculates duplex parameters on RNA-RNA thermodynamics instead of sequence-based alignment. For False-Discovery Rate correction, the genome-wide numbers of *MIRNA* loci and phasi-producing loci identified by ShortStack and PhaseTank, respectively, were used as the basis for independent tests.

**Table 1.** Schematic of various differential expression comparisons described in Results.

|  |  |  |
| --- | --- | --- |
| Experiment | Comparisons | DESeq2 Statistical test |
| Table 2. Effect of low fluence UV-B on *in vitro* plantlets | +UV-B plantlets **vs** -UV-B plantlets. Two biological replicates | Wald-Log test |
| Table 3. Effect of UV-B on berries | +UV-B (low fluence, high fluence and field samples) **vs** coordinate -UV-B (low fluence, high fluence and field samples). Three biological replicates | Wald-Log test |
| Table 5. Effect of UV-B fluence on berries | +UV-B and -UV-B high fluence **vs** co-ordinate +UV-B and -UV-B low fluence. Four biological replicates | Wald-Log test |
| Table 6. sRNAs differentially expressed during berry development | -3 WAV, 0, and +3 **vs** +6 WAV coordinate to three conditions (low fluence, high fluence and field samples) and two treatments (+UV, -UV). Three biological replicates | Likelihood Ratio Test |

### 2.4 Identification of conserved miRNAs and targets

To identify the conserved miRNAs, all miRNA candidates and sRNA clusters called by ShortStack were annotated to genome coordinates of 163 vvi-miRNAs deposited in miRBase (version 21; Kozomara & Griffiths-Jones, 2014; http://www.mirbase.org/). To identify the mRNA targets that are sliced by miRNAs, CleaveLand 4.4 (Addo-Quaye *et al*., 2009) analysis was performed. Four degradome datasets were used for this analysis. Publicly available grape degradome data (Pantaleo *et al*., 2010; GSM458931) was pre-filtered through *Vitis vinifera* ncRNAs in Rfam (Kalvari *et al*., 2017) (Suppl. Table. 1b) and was used as the *bona fide* degradome input. We also created three independent “pseudo-degradome” libraries by concatenating small RNA libraries. The rationale for using small RNA libraries as pseudo-degradome inputs for CleaveLand analyses is grounded in our interest to study those 5′-monophosphate sliced polyadenylated transcripts which are templates for production of phasiRNAs triggered by miRNA activities. Such species of 21 nt diced dsRNA products are well-represented in sRNA libraries (Luo et al., 2009; Rock, 2013). Further, independent corroboration of slicing activities from different sources improves the statistically defensible rigor of CleaveLand (Addo-Quaye *et al*., 2009). We therefore concatenated all 28 UV-B test sRNA libraries and employed them as a biological replicate, as well as the four sRNA libraries from Pantaleo et al. (2010) and independently the 14 sRNA datasets used for *MIRNA* empirical characterizations with ShortStack (Belli Kullan *et al*. 2015; Paim-Pinto *et al*. 2016; Sun *et al*. 2015; Wang *et al*. 2012, 2014; and Zhai *et al*. 2011). The identification of most of the canonical, deeply conservedmiRNA targets validated the robustness of the approach. For example, Pantaleo *et al*. (2010) identified targets for 13 conserved miRNA families out of 24, but not miR168, miR395, miR396 and miR403 despite high effector abundances. We validated 126 targets from 27 of 31 deeply conserved miRNA families, including for miR168, −395, −396, and −403 (Supplementary Table 2a and Supplementary File S4). When a known canonical miRNA target was not validated due to absence of slice signature reads in degradome datasets (miR397, −399, −408, and −827), we relied on Generic Small RNA-Transcriptome Aligner (GSTAr) output with low Allen scores to identify the missing canonical predicted targets (all found except for miR827 targets; Supplementary Table 2a). It is noted that vvi-miR397 target *VIT_08s0040g01790/GSVIVT01025694001/LACCASE11* has been previously validated by 5′ RNA Ligase-Mediated Rapid Amplification of cDNA Ends (RLM-RACE) in grapevine (Sun *et al*., 2015). We further obtained evidence for novel targets of some conserved miRNAs (miR169, −395, −396, −477, −530, −535, and −828) and for grapevine-specific miRNAs miR3623, −3624, −3626, −3629, −3631, −3632, 3633, and miR3635 (Supplementary Table 2a, Supplementary File S4). The *JAZ4* homologue of miR169 novel target *VIT_01s0011g05560/JAZ3_1* transcription corepressor (Supplementary File S4, p. 14) has recently been demonstrated along with canonical miR169 targets *NF-YAs* to be an important effector of temperature adaptation and phenology in Arabidopsis by interaction with APETELA2-like TARGET OF EAT1/2 (targets of miR172; Supplementary File S4, p. 17; Supplementary Table 2a) that negatively regulate FLOWERING TIME (Gyula *et al*., 2018). The miRNA targets were also predicted in parallel using the psRNATarget: A Plant sRNA Target Analysis Server (Dai *et al*., unpublished) and Plant Non-coding RNA Database (PNRD) (Yi, Zhang, Ling, Xu, & Su, 2015). The expectation threshold of psRNATarget was set to an Allen score = 4 (Fahlgren & Carrington, 2010) for target predictions (Supplementary Table 2b).

### 2.5 RNA blotting and Densitometry

Total RNA (15 μg) was loaded onto a 17 % polyacrylamide gel with 7M urea and was electrophoresed in 0.5X TBE at 200 V. RNA was blotted onto the positively charged Amersham Hybond-N^+^ nylon membrane (GE Healthcare Life Sciences, USA) using the Owl semidry transfer apparatus (Thermoscientific, USA) and the membrane was UV-crosslinked (SpectroLinker XL-1500, Spectroline, Westbury NY). A 22-nt anti-vvi-miR828 Locked-Nucleic Acid oligonucleotide (Valoczi *et al*., 2004) (Exiqon Inc., Woburn, MA; www.exiqon.com) was end labelled with [γ-^32^P]ATP, 6,000 Ci/mmol, (Perkin Elmer, www.perkinelmer.com) and used as probe. Hybridization was performed at 37° C for 16-20 h with PerfectHyb™ Plus hybridization buffer (Sigma). Post-hybridization washes were performed as follows: The hybridization solution was discarded, and the blots were washed sequentially with 2X SSC/0.2 % SDS, 0.5X SSC/0.2 % SDS and 0.1X SSC/0.2 % SDS. Each wash was done for 30 min at 37° C in the hybridization oven. As an equal loading control, the membrane blot was re-probed with end labelled [γ-^32^P]ATP 5S rRNA oligonucleotide. RNA blots were scanned using a Storm 860 PhosphorImager and sRNA signals were quantified using the ImageQuant TL software (v2003, GE Healthcare). We segmented the miR828, 5S rRNA band areas into seven vertical subsections of equal area per lane. Values of five subsections were averaged after discarding the leftmost and rightmost subsections which in some lanes showed gel migration artifacts. An area of the same subsection dimensions adjacent to bands was subtracted from the signal to remove background noise. A ratio of miR828 average signal divided by the control 5S RNA average signal from each lane was calculated to normalize across samples for lane loading variations. The test comparison (UV treatment -fold change) for expression strength was calculated as the ratio of normalized plus UV sample signal divided by the cognate minus-UV sample normalized signal (set to unity) for a given tissue (*in vitro* plantlet replicates), developmental time point, greenhouse fluence condition, and field condition.

### 2.6 Quantitative Real-Time PCR

cDNA amplification was conducted as in Matus *et al*. (2009). Quantification of relative gene expression was carried out as in Dauelsberg *et al*. (2011), following a thermocycling program of 95 °C for 10 min followed by 40 cycles at 95 °C for 20 s, 55-60 °C for 20 s, and 72 °C for 20s, followed by 71 cycles increasing from 50 to 96 °C at increments of 0.5°C per cycle for 30s to obtain a melting curve. Standard quantification curves with serial dilutions of PCR products were constructed for each gene to calculate amplification efficiency. Melting temperatures, primer sequences and amplification efficiency coefficients are shown in Supplementary File S7. *VviUBIQUITIN1* (VIT_16s0098g01190) was used as reference gene for calibration. To differentiate the transcripts of *Tas4a, Tas4b* and *Tas4c* and to amplify specific transcripts, primers were designed flanking the endonucleolytic slicing sites for miR828 and TAS4 3′D4(-) in highly variable regions (Supplementary Fig. S2A). qRT-PCR amplicons were run in 1% agarose gels by electrophoresis and purified and sequenced to validate appropriate designs. All experiments were performed with three (greenhouse experiment) and four (field experiment) biological replicates and two technical replicates.

### 2.7 Data Availability

The raw data files of UV sample libraries can be accessed from the National Center for Biotechnology Information (NCBI) under BioProject ID PRJNA431619.

## 3 Results

sRNA libraries were constructed for four *in vitro*-grown leaf samples (control and UV-B treated, two biological replicates) and 24 berry skin samples and sequenced by Illumina NextSeq500v2 (1 x 75 x 6 cycles for indexes). We obtained a total of 356,090,480 raw reads (Supplementary Table 1a). After adaptor trimming and removal of rRNA, tRNA, snRNA, and TE-like sequences longer than 18 nt found in the 28 libraries (Supplementary Table 1a), there remained 231,853,790 clean reads (65.7% of raw reads), of which 8% on average were unique within libraries and 5.2% were unique across all libraries. Reads longer than 27 nt were retained in the dataset for quality control assessment of libraries, which contained degraded grapevine mRNAs and 18S and 25S rRNA species of discreet 32 and 48 nt sizes not filtered with Arabidopsis rRNA sequences, which were filtered out (data not shown). ShortStack mapped 31% of reads uniquely to the reference genome, 32.8% of reads mapped to multiple loci and were binned proportionally based on flanking sequence uniqueness (Johnson *et al*., 2016), and only 0.3% of reads mapped to more than 50 loci and were discarded. Principal component analysis of berry samples showed a near majority of variation in abundances of sRNAs mapping to 40,678 loci between libraries was strongly associated with berry development (48%, PC1) (Fig. 1). The chronological sequence from green berry (−3 WAV), veraison, +3 WAV, and +6 WAV berry skin clustered coordinately and continuously irrespective of samples being collected from greenhouse or field (Fig. 1).

**Figure 1.**
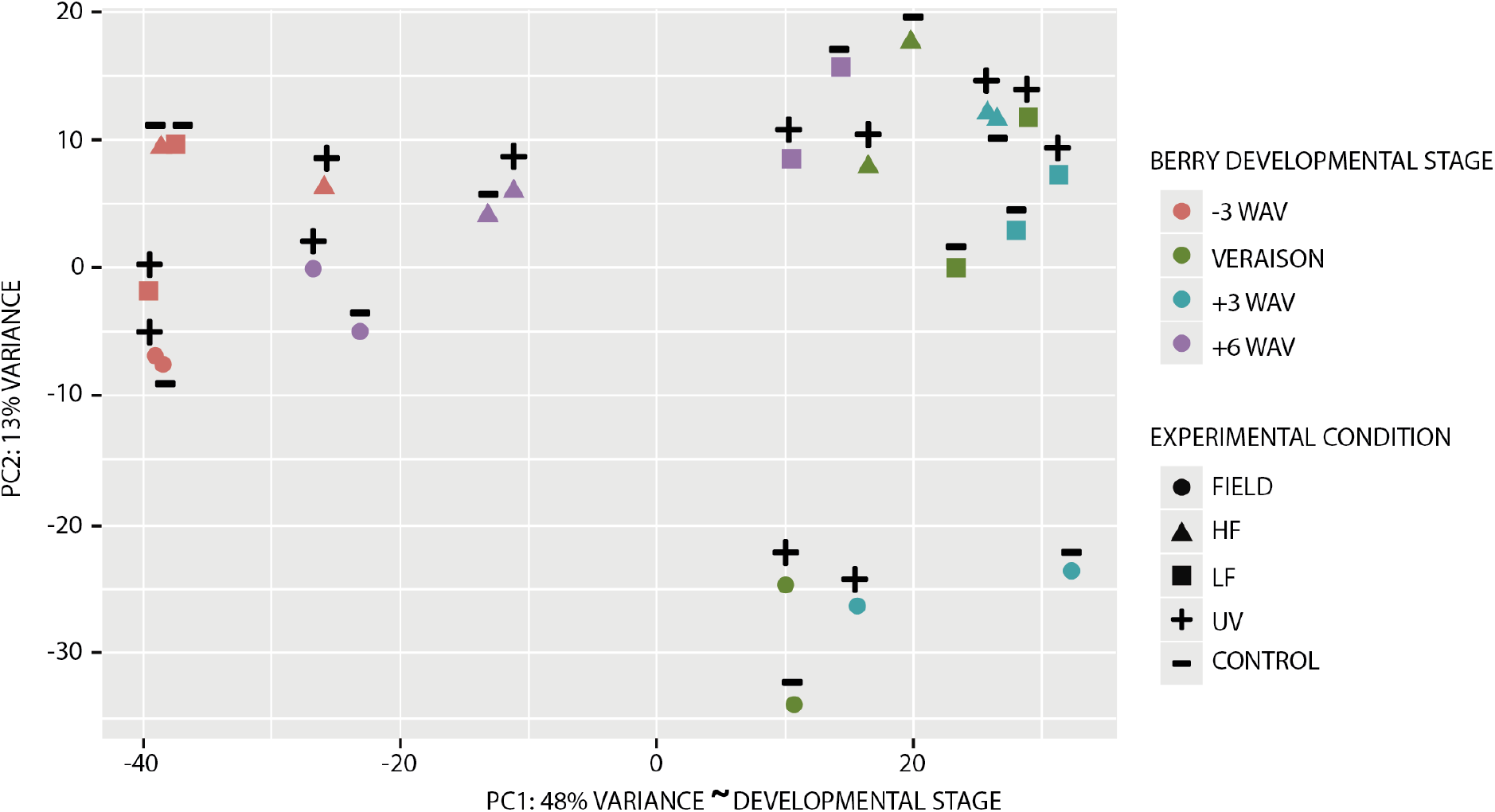
Principal component (PC) analysis of all 24 grape berry library sRNA-generating loci subjected to differential expression analysis. The percentage of variation is depicted in the PC1 and PC2 axes. Based on temporal clustering of colors, PC1 is inferred to capture the developmental stage (WAV, weeks after veraison) of samples. HF: high-fluence UV-B greenhouse berries; LF: low-fluence UV-B greenhouse berries.

The size distributions of trimmed reads were analyzed. All libraries had distinct peak abundances at 21- and 24-nt (Supplementary Fig. S1). Consistent with previous reports in grapevine (Pantaleo *et al*., 2010; Paim-Pinto *et al*., 2016), the 21-nt species was more abundant when compared to the 24-nt sRNA species. To identify the conserved miRNAs, all miRNA candidates and sRNA clusters called by ShortStack were annotated to genome coordinates deposited in miRBase. The following *MIRNAs* were not found in the ShortStack dataset: *vvi-MIR169akw*, −171dgj, −395k, −399f, −535a, −845ade, and *vvi-MIR3631c*. Other *MIRNAs* identified in the dataset and reported in the literature but not in miRBase were vvi-miR529, vvi-miR530 (Pantaleo *et al*. 2010), vvi-miR827 (Alabi, Zheng, Jagadeeswaran, Sunkar, & Naidu, 2012; Han *et al*., 2016; Pantaleo *et al*. 2010), vvi-miR858 (Pantaleo *et al*., 2010), vvi-miR4376, vvi-miR5225 (Wang *et al*., 2011; Xia *et al*., 2013), and vvi-miR7122 (Pantaleo *et al*., 2016; Xia *et al*., 2013). Also identified in the dataset as differentially regulated were 13 novel candidate *MIRNAs*, including several with weak hairpin homology to other plant species in miRBase, e.g. miR5741/8685, miR7099, and miR8761. The novel *MIRNA* hairpin sequences, refseq genome coordinates, and read counts are documented in Supplementary File S6. The miRNA expression profiles under different experimental set up and their experimentally confirmed/putative mRNA targets are discussed in detail below.

### 3.1 UV-B responsive miRNAs in *in vitro-grown* plantlets

Based on the observation that UV-B irradiation increased *VviHY5* co-expressed genes and favored flavonol (and potentially anthocyanin) accumulation in leaves of *in vitro* grown- and treated plantlets (Loyola *et al*., 2016: Matus *et al*., 2017), and to provide perspective in defining berry-specific relationships relative to leaf UV-B (Cavallini *et al*., 2015) sRNA responses, two biological replicates irradiated with low fluence UV-B and two plants with no UV-B treatment were analyzed for differential expression of miRNAs (Table 2). Two UV-B responsive miRNAs, vvi-MIR5225 and vvi-MIR160e, the former targeting phasi-RNA spawning locus *Ca*^2+^-*ATPase10* (Supplementary Table 2a) and latter targeting ARF10/17 TFs (Supplementary Table 2a; 2b; Supplementary File S4, p5) were identified as significantly up-regulated (Table 2). miR5225 is a 22 nt species evolutionarily related to miR3627 and miR4376 (Xia *et al*., 2013; Wang *et al*., 2011). In grape miR5225, miR3627 and miR4376 target the 5′UTR intron2 region of calcium-ATPase10/VIT_11s0052g00320, 14-40 nt upstream of the translation start site independently demonstrated by the Cleaveland analyses of miR4376 and iso-miR3627 in public and Pantaleo *et al*. (2010)sRNA pseudo-degradome datasets (Supplementary Table 2a; Supplementary File. S4, pp. 56, 57). The orthologous genes in cassava, hop, and mangrove have expressed sequence tags (NCBI JG970650.1, ES653410.1, DB992577.1) mapping to the miR5225/3627/4376 target site and downstream through the translation start site, supporting the claim that *VIT_11s0052g00320* intron2 is mis-annotated.

**Table 2.** Differential expression of miRNAs in in vitro plantlets subjected to UV-B radiation. C: Conserved miRNA; NC: Non-1399 conserved miRNA; N: Novel miRNA

| miRNA | log2FC | P value* | Validated Targets | Species in which targets are predicted | C/NC/ N | miRNA-triggered Phasi Locus | Short-Stack Phase Score | Reference |
| --- | --- | --- | --- | --- | --- | --- | --- | --- |
| vvi-MIR5225 | 1.67 | 0.022 | calcium-transporting ATPase10, 5' UTR intron | *Medicago truncatula*<br>*Ammopiptanthus mongolicus* | C |  | 121.3 | Xia *et al.*, 2013<br>Gao *et al.*, 2016 |
| vvi-MIR160e only 17% as abundant as star | 1.18 | 0.026 | Four Auxin Response factors ARF 10/16/17-like | Many | C |  |  | Pantaleo *et al.*, 2010<br>This report |
| vvi-MIR482 | 0.63 | 0.21 | 14 NBS-LRRs\$ | gymnosperms, dicots | C | *TAS11*;<br>NC_012019.3_chr13: 15496204-15497757^ | 1737 | This report<br>Shivaprasad *et al.*, 2012; Xia, Meyers, Liu, Beers, & Ye, 2013; Li *et al.*, 2012 |

\*Bonferroni-Hochberg Family-wise False Discovery Rate-adjusted ( $\alpha=0.1$ ) , based on all 172 *MIRNA* loci tested.
^ log2FC by UV treatment = 0.94, $P = 0.05$ . Candidate *TAS*-like locus, based on NCBI GenBank ESTs XR\_002031633.1 and XR\_787251.2 "uncharacterized ncRNA"; no homology to *sly-TAS5* (Li *et al.*, 2012). Overlaps 5' end of *VIT\_13s0047g00100*.

Although vvi-MIR482 expression was not significantly changed by UV treatment in *in vitro* plantlets (*P* = 0.21, possibly due to small sample size and high read number), PhaseTank analysis revealed a non-coding phasi-RNA candidate *TAS* locus (chr13:15496204-15497757) (ShortStack phase score 1,737) generating a tasiRNA [3′D3(-)] by vvi-miR482 cleavage (Allen score=5.0; *P* < 0.07) (Supplementary Table 4 and File S5). This novel candidate *TAS* locus significantly up-regulated by UV treatment (Table 2) was named vvi-TAS11 following the nomenclature of Zhang *et al*. (2012). vvi-TAS11 maps to the 5′ end of an unannotated 87aa peptide-encoding locus (*VIT_13s0047g00100)* with ESTs (GenBank XR_002031633.1, XR_787251.2) annotated as ‘uncharacterized non-coding RNA’. Degradome evidence (Allen score 4.0; Supplementary Table 4) supports cleavage of a phasi-RNA-producing (phase score =93.5) Leucine-Rich-Repeat (LRR) transcript *VIT_18s0001g07270* by the tasiRNA3′-D3(-), and is predicted to target another phasi-RNA-producing LRR (phase score 29.2) *VIT_13s0106g00020*, 12 NB-ARC-LRRs (Supplementary Tables 2b, 4), and the putative cation/hydrogen exchanger *Vvi-CHX15/ VIT_04s0044g01470*.

Induction of *MYBA6* and *MYBA7* in vegetative tissues upon UV-B radiation has been observed previously (Matus *et al*., 2017). Similarly, miR828/TAS4-mediated post-transcriptional regulation of *MYBA6* and *MYBA7* by production of TAS4-3′D4(-)-triggered phasi-RNAs is also well documented in leaves (Rock, 2013; Zhang *et al*., [2012] mis-annotated *TAS4* as VviTAS7). The extremely low levels of mature miR828 (~1 per 20 million reads) in the sRNA libraries, including leaves where it was expected to be maximal, made it difficult to quantify by deep sequencing. Hence, an RNA blot analysis with Locked Nucleic Acid-anti-vvi-miR828 as probe was performed to verify the expression profile of *MIR828* cluster siRNAs (miR828* was the major species found at 2.6 reads per million; Supplementary Table 3) quantified by ShortStack. As predicted, mature miR828 was maximal in leaves, but there was no obvious difference in miR828 levels in response to UV-B in plantlets (Fig. 2; Panel A; but see below).

**Figure 2.**
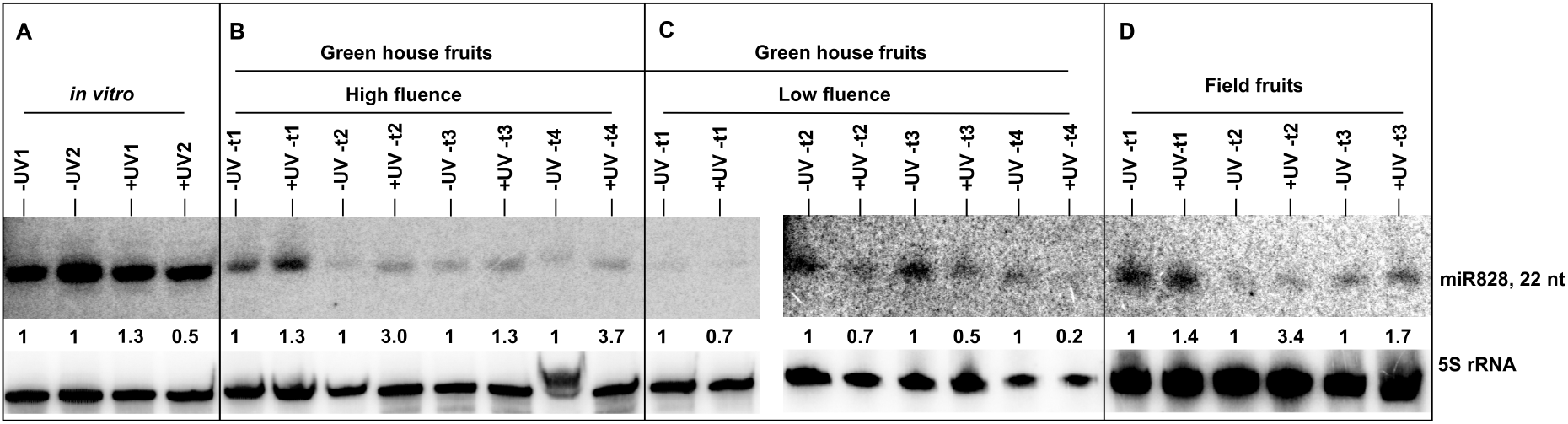
sRNA blot of miR828 abundance in UV response samples subjected to deep sequencing. The quantitation of mature miR828 abundances by densitometry was used to validate ShortStack quantitation of relative vvi-*MIR828* sRNA cluster abundances, which are used in statistical analyses as a proxy for mature miR828 species abundance since mostly miR828* species were sequenced from sRNA libraries. **A)** UV-induced expression of miR828 in *in vitro* plantlets. –UV: No UV-B-2treatment; +UV-B: 6 hr of low UV-B radiation (0.15 W m irradiance). **B)** UV-induced expression of miR828 at different stages of berry development in greenhouse conditions subjected to high fluence −2 UV-B. –UV: No UV-B treatment; +UV-B: (~ 0.3 W m^-2^ daily for 5 hr). **C)** UV-induced expression of miR828 at different stages of berry development in green house conditions when subjected to low −2 fluence UV-B. –UV: No UV-B treatment; +UV-B: (~ 0.1 W m^-2^ daily for 10 hr). **D)** UV-induced expression of miR828 at different stages of berry development in field conditions UV-B. –UV: Solar UV-B blocked by 100 μm clear polyester film; +UV-B: No filters. Panels B-D: t1, t2, t3 and t4 corresponds to Weeks −3, 0, 3, and 6 after veraison (WAV), respectively. As loading control, 5S rRNA probe was hybridized to the same membrane. The relative abundance of miR828 in test samples (+UV-B) is presented as the ratio compared to normalized abundance of -UV-B controls (set to unity).

### 3.2 UV-B responsive small RNAs in berry skins from greenhouse and field experiments

sRNA libraries were constructed from total RNA extracts of berry skins (−3, 0, +3 and +6 WAV) harvested from greenhouse and fields treated with and without UV-B light. The high- and low-UV irradiated greenhouse and field samples were classed as +UV-B test samples and corresponding minus UV-B as control samples for differential expression analysis (Table 1). Eleven UV-B responsive miRNAs were identified with statistically significant differential expression (independent of developmental time points), of which nine were up-regulated (vvi-MIR395n, vvi-MIR3627, vvi-MIR535c, vvi-MIR3624, vvi-MIR171b, vvi-MIR156f, vvi-miR4376, vvi-MIR319c) and two were overall down-regulated (vvi-MIR477b, vvi-MIR530) (Table 3). However, further analysis (described below) showed those two down-regulated *MIRNA* loci are up-regulated in response to high-fluence UV-B. The up-regulated UV-B-responsive miRNA targets were validated by CleaveLand analysis and corroborated by psRNATarget, Plant Non-coding RNA Database (PNRD) (Yi *et al*., 2015), and evidence from available literature (Table 3; Supplementary Tables 2a, 2b; Supplementary File S4). Cleaveland analysis validated Metal Ion Binding protein, previously reported by Sun *et al*. (2015) and Pagliarani *et al*. (2017) as the target of MIR3624-3p (Supplementary Table 2a; Supplementary File S4, p. 62),despite the *MIR3624* locus produces abundant phasi-RNAs from the antisense strand (Supplementary Table 3).

**Table 3.** Differential expression of miRNAs in berries subjected to UV-B radiation. ***C***: Conserved miRNA; ***NC:*** Non-conserved miRNA; ***N:*** Novel miRNA

| miRNA | log2FC | P value* | Validated Targets | Species in which targets are predicted | C/NC/N | Phasi Locus | ShortStack Phase Score | Reference |
| --- | --- | --- | --- | --- | --- | --- | --- | --- |
| vvi-MIR395n | 1.46 | 0.0001 | ATP sulfurylase 1; AST68, Low affinity sulfate transporter (miR395l,h) | Many | C |  |  | Paim-Pinto *et al.*, 2016; Sun *et al.*, 2015 |
| vvi-MIR3627-5p isomiR | 1.28 | 0.0002 | calcium-transporting ATPase10, 5' UTR intron2 | Grapevine, peach, apple | C |  | 121.3 for *MIRNA*=63 | Xia *et al.*, 2013; Gao *et al.* 2016 |
| vvi-MIR535c | 1.01 | 0.0002 | DNA primase large SU; SPLs (bulge at nt6 seed region) | Many | C |  |  | Carra *et al.*, 2009<br>Pantaleo *et al.*, 2010<br>Han *et al.*, 2016<br>Paim-Pinto *et al.*, 2016 |
| vvi-MIR3624-3p | 0.94 | 0.0002 | METAL ION BINDING PROTEIN-RELATED, Proline-rich family protein | Grapevine | NC |  | 373.3 for *MIRNA* | Sun *et al.*, 2015<br>Pagliarani *et al.*, 2017<br>this report |
| vvi-MIR171b isomiR | 0.74 | 0.006 | GRAS family scarecrow-like protein<br>SCR4-L SCR-L22 SCR-L15 | Many | C |  |  | Pantaleo *et al.*, 2010;<br>Wang *et al.*, 2013 |
| vvi-MIR477b mature is only 0.5% of star species | -0.97 | 0.02 | repC/Pol III GAMMA-TAU SU; Elongation factor 1B-gamma; EDA9, D-3-phosphoglycerate dehydrogenase | Dicots | C |  |  | Paim-Pinto *et al.*, 2016<br>This report |
| vvi-MIR530a | -0.77 | 0.025 | AGO1 (two homologues, intron and 3' UTR); Plus3 domain; PPR DYW deaminase | many | C |  |  | Alabi *et al.*, 2012; Han *et al.*, 2016; Liang *et al.*, 2013 |
| vvi-MIR156f | 0.68 | 0.028 | 9 squamosa promoter-binding-like SPL TFs | many | C |  |  | Pantaleo *et al.*, 2010; Casati *et al.*, 2013; Wang *et al.*, 2013 |
| vvi-miR4376 | 0.79 | 0.05 | calcium-transporting ATPase10 5' UTR intron2 | dicots | C |  | 121.3 | Wang *et al.*, 2011; Xia *et al.*, 2013 |
| vvi-MIR319c | 0.68 | 0.055 | TCP4-L; TCP24-L | many | C |  |  | Paim-Pinto *et al.*, 2016 |
| vvi-MIR828 | -0.48 | NA |  | Eudicots, palms | C | Vv-*TAS4b*: NC_012020.3_chr14: 21534542-21535069^ | 7,566 | Hsieh *et al.*, 2009; Luo *et al.*, 2012; Rock, 2013 |
| vvi-MIR482 | 0.70\$ | 0.018 | 10 NB-LRRs\$ | Gymnosperms, dicots | C | *sly-TAS5*-like NC_012024.3_chr18: 20665102-20666018 D12(-) → VIT_00s0238g00130; D(6+) → VIT_12s0034g01310 and VIT_12s0034g01350 | 2,456.3 | This report Li *et al.*, 2012; Xia *et al.*, 2013 |
| unknown | 0.59\$ | 0.028 | | | N | NC_012007.3_chr1: 13356141-13357440 (Phasi trigger: unannotated PPR, VIT_11s0016g02500) | 98.4 | This report |

\*Bonferroni-Hochberg Family-wise False Discovery Rate-adjusted ( $\alpha=0.1$ ) , based on all 172 *MIRNA* loci tested. NA: not analyzed due to low reads abundance. \$ LFC refers to candidate target phase locus. NB-LRR VIT12s0034g01310 (phase score=342.8), predicted target of chr18 tasiRNA D6(+) identified by PhaseTank, and VIT\_12s0034g01350 (phase score=398.3); Supplementary Table 4, File S5. Validated NB-ARC target of sly-TAS5-like tasiRNA D12(-) VIT\_00s0238g00130 by degradome data, and seven other predicted NB-ARC targets (Suppl. Tables 2b and 4; Suppl. File S5). ^ *TAS4b* LFC by UV = 1.1 ( $P = 0.005$ ; family-wise FDR-adjusted).

Cleaveland and PhaseTank analysis identified vvi-MIR482 targets a second *TAS* locus (chr18: 20665102-20666018) annotated as overlapping the 3′ UTR of an 83aa transmembrane peptide transcript *VIT_18s0072g01090* (Supplementary Table 2a, 2b; Supplementary File S4, p. 37). An abundant tasiRNA 3′D12(-) significantly up-regulated by UV is analogous by concordance of phase (but not by homology) to sly-TAS5 (Li, Orban, & Baker, 2012; data not shown) and likewise is predicted to target eight NB-ARC/TIR-LRR transcripts (Supplementary Table 2b, File S5), thus we name it vvi-slyTAS5-like. This candidate *TAS* locus produces a tasiRNA3′-D6(+) predicted to target NB-LRRs *VIT_12s0034g01310* and *VIT_12s0034g01350* (Supplementary Table 4 PhaseTank prediction, Supplementary File S5). Degradome evidence demonstrated the tasi-RNA D6(+) from vvi-slyTAS5-like also directs cleavage (Allen score 3.5) of *VIT_13s0019g03540* predicted to encode an Armadillo-type fold domain, found in nuclear cap-binding protein CBP80 and BB-LRR variants with 72% homology to regulator of nonsense transcripts UPF2/AT2G39260, and *VIT_07s0031g00350* (Allen score 3.0) encoding an S-adenosyl-methionine-dependent methyltransferase in the family of catechol/caffeoyl-CoA phenylpropanoid biosynthesis enzymes. Another PHASI locus with an unknown trigger produced an abundant siRNA up-regulated by UV-B with homology to miR6173 (Liu, Wang, Sun, Liang, & Wang, 2015; data not shown). We identified from degradome analysis a phasi-RNA-producing and *cis*-cleavage active locus triggered by miR390 possibly involving the 2_21_ two-hit mechanism (Axtell, Jan, Rajagopalan, & Bartel, 2006; Supplementary Table 2a; Supplementary File S4, p. 18; Supplementary File S5) associated with AGO7 complex (Montgomery *et al*., 2008; Rajeswaran & Pooggin, 2012). These phasi-RNAs are generated from *VIT_12s0059g01410* encoding a 66aa signal peptide with limited homology to Arabidopsis *TASIR-ARF/At5g57735* and distinct from *VviTAS3* triggered by miR390 (Xie *et al*., 2005; Zhang *et al*., 2012; Supplementary File S4, p. 19; Supplementary File S5, see below). In addition, a phasi-RNA-producing locus (unannotated, but homologous to rice Oryza|ChrUn.fgenesh.mRNA.47_GX23P; https://phytozome.jgi.doe.gov) was predicted to be targeted by miR2111 (Supplementary Table 4, prediction sheet), as well evidence for slicing of NHD1, Sodium hydrogen antiporter (Supplementary File S4, p. 58), but was not significantly differentially regulated by UV-B and not considered further.

Matus *et al*. (2017) reported berries irradiated with UV-B delayed the decrease in *MYBA5/6/7* transcripts after the onset of veraison, supporting the role of *MYBA6* and *MYBA7* in induction of pigmentation during fruit development. Luo *et al*. (2012) and Hsieh *et al*. (2009) characterized in Arabidopsis an auto-regulatory feedback loop involving up-regulation of miR828/TAS4 by AtPAP1/MYB75 (an orthologue of *VviMYBA6/7)* and being themselves targets of the negative regulators miR828/TAS4. Piya, Kihm, Rice, Baum, and Hewezi (2017) recently described a similar auto-regulatory loop for miR858 and *MYB83*. If such feedback loops are conserved in grapevine then an increase in *MYBA6/7* transcript levels would increase the levels of miR828- and/or miR828-triggered *TAS4*-3′-D4(-) from *VviTAS4abc* loci, which cleave *MYBA6/7* differentially to generate phasi-RNAs that amplify post-transcriptional silencing of *MYBA6/7* (Rock, 2013). Inference of miR828 expression by deep sequencing read abundances of miR828*, which was much more abundant than the rarely sequenced miR828, suggested overall down-regulation of *MIR828* locus by UV-B treatment (Table 3). Yet we observed significant UV-B induced accumulation of *TAS4b-* 3′D4(-) (Table 3 ^footnote; Supplemental Table 3) which is functionally contingent upon miR828 activity, consistent with the results of Matus *et al*. (2017) and the hypothesized auto-regulation of miR828/VvTAS4abc loci. This hypothesis was substantiated by CleaveLand analysis of sRNA pseudo-degradome in which we observed miR828 slicing of *VviTAS4a/b/c* and the phased *TAS4ab-* 3′D4(-) species targeting *MYBA7, MYBA6*, and *MYBA5/MYB113-like/VIT_14s0006g01340* (Supplementary Table 2a; File S4, pp. 38-43). Because *vvi-MIR828* was not significantly correlated to UV-B by DESeq2 analysis across all fluence and field treatments, we characterized mature miR828 expression alone by RNA blot (Fig. 2). Interestingly, semi-quantitative densitometry analysis of band intensities showed reproducible up-regulation of miR828 by high fluence UV-B treatments in both greenhouse and field, and down-regulation of miR828 by low fluence UV-B greenhouse treatments (Fig. 2B-D), suggesting a complex regulation (up-regulation by high UV fluence and down-regulation by low UV fluence) of *vvi-MIR828* locus by UV-B.

The functional significance of differentially expressed miRNAs upon high UV-B influence in the greenhouse was tested by real-time quantitative PCR (qRT-PCR) of the expression levels of validated target genes from the same samples used to construct the sRNA libraries. The results are shown in Fig. 3. An inverse correlation was found between the observed miRNA dynamic and each target gene expression profile during berry development. Differentially expressed miR156f, miR3632-3p (homologous to miR482) and miR530a throughout berry development displayed an inverse relationship with their cognate target genes *Squamosa promoter binding protein-like 2 (SPL2_3/VIT_11s0065g00170)* (Supplementary Table 2a; Supplementary File S4, p. 1), numerous homologues of *VIT_13s0067g00790* encoding Leucine-rich repeat protein (LRR) (Supplementary Table S2b; Supplementary File S4, p. 66), and *Plus3 domain* protein-coding genes (Supplementary Table 2a; Supplementary File S4, p. 11), respectively (Fig. 3A-C). A direct relationship was observed between miR482 and derivative tasi-RNA *TAS11* 3′-D3(-) abundance changes during berry development and in response to high UV-B (Fig. 3D, left and middle panels). Likewise to miRNA:target dynamics, the *TAS11* regulatory cascade showed evidence of activity against downstream target NB-ARC LRR protein gene (Fig. 3D, compare middle and right panels). Interestingly, a direct concordance was observed between the *TAS4c* 3′D4(-) tasi-RNA and its primary (undiced) *TAS4c* transcript (Fig. 3D), supporting the notion that miR828 induction by high fluence UV-B (Fig. 2B, D) during berry development results in increased *TAS4* expression marked by accumulation of *TAS4* phasi-RNAs triggered by miR828 (Fig. 3E, left panel; Supplementary Table 2a; File S4, pp. 38-40). Similar results were observed between *TAS4a* 3′D1(+), *TAS4b* 3′D4(-), and *TAS4c* 3′D4(-) tasi-RNAs and their respective primary *TAS4* transcripts from high-fluence UV-B field samples (Supplementary Fig. S2B-D). However, these findings raise the question of why *TAS4* primary transcript relative abundances are not *reduced* by UV-B treatment; i.e. an inverse relationship with miR828 (Fig.2) as seen for other miRNA targets and tasi-RNA cascades (above). We observed high fluence UV-B treatments in greenhouse and field significantly delayed the down-regulation of *MYBA6* and *MYBA7* at the latter stages of berry development (Supplementary Fig. S2E, F right panels; Matus *et al*., 2017). We propose that the observed concordant expressions of miR828 (Fig. 2), *TAS4abc* primary transcripts and diced tasi-RNAs (Fig. 3E, Supplementary Fig. S2B-D) and the derived *MYBA6* and *MYBA7* phasi-RNAs spawned by the activity of TAS4ab-3′D4(-) (Supplementary Fig. S2E, F left panels; Supplementary File S4, pp. 41, 42) are strong evidence in support of a functional orthologous *MYBA6/7* auto-regulatory loop as described in Arabidopsis (Hsieh *et al*., 2009; Luo *et al*., 2012) acting either directly or indirectly to activate expression of both *MIR828* and *TAS4abc*.

**Figure 3.**
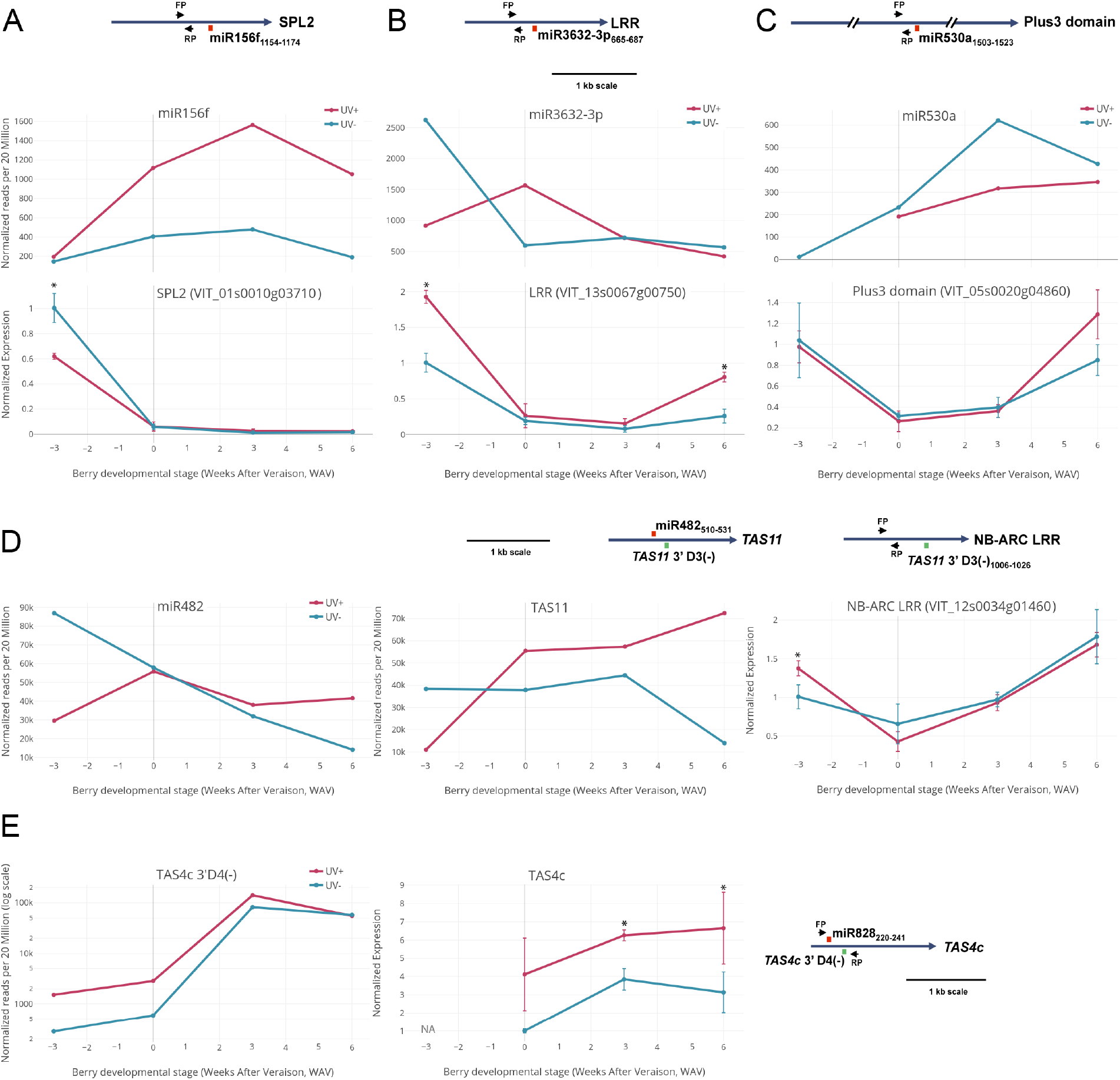
Functional validation of differentially expressed miRNA activities under high fluence UV-B by qRT-PCR of target genes. **Left Panel A-D:** miRNA/siRNA deep-sequencing profiles at different berry developmental stages (weeks after veraison, WAV) under high UV-B fluence in greenhouse. **Right Panels A-D:** miRNA target gene expression profile at different berry developmental stages under high UV-B fluence. **A)** miR156f and its target Squamosa promoter binding protein-like 2 (SPL2) expression profile. **B)** miR3632-3p and its target Leucine-rich repeat protein (LRR) expression profile. **C)** miR530a and its target Plus3 domain protein expression profile. **D)** miR482 expression profile (left panel). **Middle Panel:** *TAS11* 3′ D3(-) *trans* acting siRNA expression profile triggered by miR482. **Right Panel:** Expression profile of NB-ARC LRR targeted by *TAS11* 3′ D3(-) tasiRNA. **E)** *TAS4c* 3′ D4(-) *trans* acting siRNA expression profile (left panel) and the expression profile of *TAS4c* undiced primary transcript (right panel; “NA”: not available. Test samples were normalized to 0 WAV minus UV-B [set to unity]). Error bars are s.d. Asterisks (*) denote significant differences based on analysis of variance (n=3 biological replicates) and comparisons using the Tukey-Kramer honestly significance test (HSD; *p* < 0.05). **Insets, A-E:** Target-miRNA/siRNA binding positions (mRNA nucleotide coordinates) and primer binding sites (arrows) are depicted to scale. Target:miRNA binding depicted in Red; Target:tasiRNA binding depicted in Green;

The observed complex regulation of miR828 by UV-B detected by RNA blot prompted us to examine other significantly differentially expressed *MIRNAs* (all analyzed by DESeq2) for a pattern of up-regulation by high UV fluence and down-regulation by low UV fluence. Table 4 lists those significant UV-B differentially regulated miRNAs and siRNAs mimicking the pattern (up in high fluence, down in low fluence) observed by RNA blot for miR828 (Fig. 2). We further endeavored to impute supporting evidence for miRNA function by meta-analysis with respect to the observed UV-B changes of miRNA abundances with co-expression analyses of validated mRNA targets using three published genome-wide transcriptomics datasets of grapevine berry skin tissues in response to 1 hr of UV-C exposure (Suzuki *et al*., 2015), five weeks exposure to UV-B after veraison (Carbonell-Bejerano *et al*., 2014), and from five red-skinned varieties sampled across berry developmental stages (Massonnet *et al*., 2017). It is noted that UV-C lamps have about 5% of their energy emitted in the UV-B band, supporting our view that these comparisons are worthwhile. Table 4 shows the normalized abundances (ratio of +UV/Control) of selected miRNAs for four time points and three environments in response to UV-B (minus UV-B set to unity), and the reported fold-change effects on validated miRNA target mRNAs (Supplementary File S4; Supplementary Table 2b) after 1 hr UV-C and five weeks UV-B treatments (UV-C treatment can be a proxy test for high fluence UV-B given that UV-C light source GL-15 lamps have power emission in the UV-B range; http://www.nelt.co.jp/english/products/safl/). Binomial distribution tests of significance for the observed inverse relationships to occur by chance (a function of number of targets) revealed significant results collectively, and individually for all 10 predicted SPL targets of miR156f, 12 of 14 LRR targets of miR482-triggered *TAS11* 3′D3(-), and four of five validated MYB targets of miR828 (*VIT_14s0066g01220, VIT_09s0002g01380, VIT_00s0341g00050*, and *VIT_17s0000g08480;* Supplementary Table 2b; Supplementary File S4, pp. 44-48). Taken together these results support that high-fluence UV-B induces miRNA accumulations that consequently effect decreased berry skin target gene expressions.

**Table 4.** Co-expression analysis of high fluence UV-B induction of miRNAs/phasi-RNAs abundances during berry development which are inversely correlated with predicted target mRNA expressions in the same skin tissues in response to UV-C pre-veraison and UV-B five weeks post-veraison from independent experiments (Suzuki *et al*., 2015; Carbonell-Bejerano *et al*., 2014).

|  |  |  |  |  |  |  |  |  |  |  |  |  |  |  |  |  |  |
| --- | --- | --- | --- | --- | --- | --- | --- | --- | --- | --- | --- | --- | --- | --- | --- | --- | --- |
|  |  | UV-B Fold Change* (green= **up**; red = **down**) |  |  |  |  |  |  |  |  |  |  |  | Inverse correlation of mRNA target expression versus miRNA/siRNA? |  |  |  |
|  |  | High Fluence Greenhouse |  |  |  | Low Fluence Greenhouse |  |  |  | High Fluence Field |  |  |  |  |  |  |  |
| miRNA/siRNA | LFC, UV-B | -3 WAV | Verai -son | + 3 WAV | +6 WAV | -3 WAV | Verai -son | + 3 WAV | +6 WAV | -3 WAV | Verai -son | + 3 WAV | +6 WAV | Validated Targets* (n; annotation) | FC, UV-C<sup>§</sup> | FC, UV-B<sup>†</sup> | FC *P*-val<sup>^</sup> |
| miR156f | 0.68 | 1.332 | 2.755 | 3.266 | 5.520 | 0.644 | 10.25 | 1.863 | 0.567 | 1.440 | 2.149 | 1.144 | 1.430 | 11; SPB | **0.75** | 0.89 | 0.02 |
| miR530b | -0.77<sup>+</sup> | n.d. | 2.556 | 0.488 | 1.183 | 0.258 | 0.853 | 0.200 | 0.065 | n.d. | 5.908 | 5.853 | 6.316 | 2; Plus3, DYW | **0.80** | 0.86 | 0.09 |
| miR477c | -0.32<sup>+</sup> | 0.010 | 3.253 | 3.129 | 2.772 | 0.017 | 0.104 | 0.116 | 0.003 | 3.082 | 1.177 | 3.039 | 0.374 | 2; GRAS-domain | n.d. | 0.84 | 0.51 |
| TAS4a, miR828 targ | 0.32 | 0.561 | 3.540 | 5.119 | 1.249 | 0.102 | 2.155 | 0.554 | 0.357 | 2.772 | 2.602 | 1.961 | 1.192 | 2; MYBA6/A7 | **0.63** | 0.97 | 0.30 |
| TAS4b, miR828 targ | 1.10 | 0.089 | 1.704 | 2.115 | 60.16 | 1.006 | 2.713 | 2.562 | 0.572 | 4.956 | 7.022 | 6.213 | 2.953 |  |  |  |  |
| TAS4c, miR828 targ | 0.14 | 5.381 | 4.825 | 1.712 | 0.955 | 0.180 | 1.348 | 1.342 | 0.513 | 0.318 | 1.777 | 1.953 | 0.750 |  |  |  |  |
| miR828targ MYB, phasi | -0.29<sup>+</sup> | 0.218 | 1.425 | 1.023 | 1.725 | 0.130 | 2.948 | 1.260 | 0.377 | 1.151 | 1.367 | 0.569 | 1.316 | 5; MYBs | **0.70** | 0.87 | 0.04 |
| miR482 | 0.07 | 0.340 | 0.965 | 1.188 | 2.936 | 0.393 | 2.583 | 2.840 | 0.624 | 1.450 | 1.678 | 0.774 | 0.855 | 2;TAS11,slyTAS5-L | **0.80** | 0.96 | 0.09 |
| TAS11-D3(-) miR482 targ | 0.17 | 0.287 | 1.467 | 1.290 | 5.191 | 0.515 | 5.015 | 1.697 | 0.419 | 1.673 | 1.167 | 0.733 | 1.279 | 14; LRRs | **0.84** | 0.86 | 0.02 |
| miR403a-e average | -0.19 | 0.504 | 1.206 | 0.205 | 1.212 | 2.764 | 2.238 | 0.347 | 0.382 | 1.566 | 0.987 | 0.678 | 2.044 | 1; AGO2 | 1.1 | 1.07 | 0.17 |
|  |  |  |  |  |  |  |  |  |  |  |  |  |  | Overall 38 targets, two experiments |  |  | 5.3E-6 |
| <sup>&</sup> Normalized to 20M reads (+1 if zero reads). Denominator is corresponding normalized -UV-B sample (unity). n.d.: not detected<br><sup>+</sup> sign negative overall because low UV-B fluence effect dominant, or low read counts independently filtered out by PhaseTank<br><sup>*</sup> See Supplementary Table 2a.<br><sup>§</sup> Data for 1 hr FC by UV-C treatment of pre-veraison berries from Suzuki *et al.* (2015). If called significantly different expression in response to UV-C, in **bold**.<br><sup>^</sup> Binomial distribution probability for FCs of al target mRNA responses to UV-C and UV-B inverse to miRNA/siRNA (up) under high fluence UV-B<br><sup>†</sup> Data for FC by UV-B treatment on 26 °Brix (ripe) berries irradiated for five weeks post-veraison from Carbonell-Bejerano *et al.* (2014). For miR403 target AGO2, includes 23°Brix sample (3 independent tests for significance). |  |  |  |  |  |  |  |  |  |  |  |  |  |  |  |  |  |

### 3.3 UV-B regulated miRNAs with different fluence-rate responses

Based on the apparent coordination for miRNA expressions by high-versus low fluence UV-B treatments, the high- and low fluence-irradiated greenhouse berry samples were tested against each other (high/low ratio across developmental time, including across controls minus UV). The field UV-B irradiated berries were not included in the test since the environmental parameters were not controlled. Table 5 lists 18 miRNAs with significant differential expression by high fluence UV-B treatment per se, of which 11 were up-regulated and seven were down-regulated. Three novel miRNAs (called as valid *MIRNAs* or missing only evidence for the star species by ShortStack and folding into a stable and unbulged hairpin) were identified to be differentially expressed in response to high fluence UV-B. We found multiple candidate family members of *vvi-MIR477* of which miRBase has annotated only two: miR477ab. The five others which respond differentially to UV-B fluence listed in Table 5 are presumed to be encompassed by the nine vvi-miRC477c,i-p candidate family members described by Paim-Pinto *et al*. (2016). Similar to the UV-B response results across greenhouse and field berry samples (Table 3), *vvi-MIR3627* was highly up-regulated under high fluence (Table 5). *vvi-MIR3627* is an atypical *MIRNA* locus in that it produces phased sRNAs and is not stranded (it produces abundant antisense transcripts (Supplementary Table 3). vvi-MIR399i, which is predicted to target phosphate transporters (Supplementary Table 2a) (Mica *et al*., 2010), vvi-MIR167c targeting *ARF* TFs (Supplementary Table 2a; Supplementary File S4, p. 9) were some of the miRNAs up-regulated in high fluence.

**Table 5.** Differential expression of miRNAs in berries subjected to high versus low fluence UV-B radiation. ***C:*** Conserved miRNA; ***NC:*** Non-conserved miRNA; ***N:*** Novel miRNA

| miRNA | log2FC | P value* | Validated Targets | Species in which targets are predicted | C/NC/N | Phasi Locus | Phase Score | Reference |
| --- | --- | --- | --- | --- | --- | --- | --- | --- |
| vvi-MIR3627 | 2.02 | 0.000005 | calcium-transporting ATPase10 | Grapevine<br>peach, apple | C |  | 121.3<br>for<br>MIRNA=63 | Wang *et al.*, 2011;<br>Xia *et al.*, 2013 |
| vvi-MIR399i | 2.10 | 0.0001 | Major Facilitator<br>PHT1_3/PHO84; inorganic<br>phosphate transporter | many | C |  |  | Mica *et al.*, 2010 |
| vvi-MIR535c | -1.19 | 0.0007 | DNA primase large SU; SPLs<br>(bulge at nt6 seed region) | many | C |  |  | Pantaleo *et al.*, 2010<br>Paim-Pinto *et al.*, 2016 |
| vvi-MIR167c | 1.37 | 0.001 | ARF6, ARF8, ARF6-Like;<br>predicted bulge target nt10 | many | C |  |  | Paim-Pinto *et al.*, 2016;<br>Wang *et al.*, 2013 |
| vvi-MIR403f | -0.81 | 0.008 | AGO2 | dicots | C |  | 2.4 | Pantaleo *et al.*, 2010 |
| vvi-MIR403d | -0.78 | 0.01 | AGO2 | dicots | C |  |  | Pantaleo *et al.*, 2010 |
| vvi-MIR3632-3p; 22 nt<br>species | 0.82 | 0.01 | six NB-ARC disease resistance<br>LRRs rga4-like | Grapevine | NC |  |  | Pantaleo *et al.*, 2016<br>Sun *et al.* 2015<br>Fan *et al.* 2015<br>Paim-Pinto *et al.*, 2016 |
| vvi-MIR169c-5p isoMIR | 1.43 | 0.013 | Five Nuclear transcription factor<br>Y subunit A- related; |  |  |  |  | Gyula *et al.*, 2018 |
|  |  |  | JAZ3_1/VvJAZ1_TIFY transcription corepressor; Zinc finger At1g75340-like | many | C |  |  | Paim-Pinto *et al.*, 2016 |
| vvi-MIR403a | -0.78 | 0.013 | AGO2 | dicots | C |  |  | Pantaleo *et al.*, 2010 |
| vvi-MIR477b | 1.5 | 0.016 | See Table 3; | dicots | C |  |  | Paim-Pinto *et al.*, 2016 |
| novel-1; 24 nt species | 1.16 | 0.026 | None predicted |  | N |  |  | This report |
| vvi-MIR403e | -0.74 | 0.027 | AGO2 | dicots | C |  |  | Pantaleo *et al.*, 2010 |
| novel-2; 24 nt species | 1 | 0.03 | None predicted |  | N |  |  | This report |
| novel_aly-MIR3435-hp-like | -0.98 | 0.04 | None predicted |  | N |  |  | This report |
| vvi-MIR403c | -0.77 | 0.046 | AGO2 | dicots | C |  |  | Pantaleo *et al.*, 2010 |
| vvi-MIR172b | 1.20 | 0.05 | Four APETALA2-like TFs | many | C |  |  | Pantaleo *et al.*, 2010 |
| vvi-MIR828 targeted *TAS4a* | 0.41 | 0.03 | MYBA6/MYBA7 | eudicots | C | NC_012020.3_chr14:21607803-21608825 | 18,631 | Rock, 2013 |
| *TAS4c* | -1.18 | 0.09 |  | eudicots | C | NC_012007.3_chr1:2960325-2961696 | 3,375.4 | Rock, 2013 |
| vvi-MIR7122; 22 nt species | 0.61\$ | 0.06 | None predicted | dicots | C | | 5,366 | Xia *et al.*, 2013 |
| vvi-MIR828 MYB target | 0.99\$ | 0.006 | MYB | Gymnosperms, eudicots | C | NC_012015.3_chr9:1161752-1163111; VIT_09s0002g01380 | 1,260.5 | Pantaleo *et al.*, 2010; Luo *et al.*, 2012 |
| vvi-MIR482 LRR target | 0.74\$ | 0.03 | NB-LRR | grapevine | N | *vvi-sly-TAS5-like*<br>NC_012024.3_chr18:20665102-20666018 D6(+) → VIT_12s0034g01350 | 398.3 | This report<br>Li *et al.*, 2012; Xia *et al.*, |
|  |  |  |  |  |  |  |  | 2013 |
| *TAS3*, miR390 target | 1.47 | 0.075 | D7(+): ARF2,3,4<br>D2(-):3' UTR of<br>TOPOISOMERASE II-ASSOCIATED PROTEIN<br>D5(-):3' UTR of Zinc finger<br>RING/FYVE/PHD | many | C | NC_012011.3_chr5: 358508-360140 | 3172.2 | Xia *et al.*, 2013;<br>Zhang *et al.*, 2012 |
| Unknown trigger | 1.47 | 0.08 |  |  |  | NC_012011.3_chr5: 6017171-6018107 | 504.6 | This report |
| *Bonferroni-Hochberg Family-wise False Discovery Rate-adjusted ($\alpha=0.1$) , based on all 189 *MIRNA* loci tested, minus those (108) loci that were independently filtered out by DESeq2 due to low counts cutoff (<7.8 base mean reads per library).<br>\$ LFC and *P* values for target LLR and MYB phasi-RNAs; degradome validated (Supplementary Table 4, File S5). | | | | | | | | |

Two interesting observations were the coordinated downregulation by high fluence UV-B of all six miR403 family members (five of them significantly), that target *AGO2* in dicots (Supplementary Tables 2a, 2b; Supplementary File S4, p. 34), which is correspondingly up regulated by UV-C and UV-B (Table 4) and associated with stress responses including DNA damage (Fátyol, Ludman, & Burgyán, 2016; Wei *et al*., 2011). The up-regulation of *MIR403f* primary transcript during berry maturation in field samples (Supplementary Fig. S3) validated our observation of significant up-regulation across developmental time points for miR403 species (Table 6). *TAS4a* targeted by vvi-miR828 (phase Score: 18,631; Supplementary Table 2a; File S4, p. 38), a miR828-targeted MYB (phase Score: 1,260, Supplementary File S4, p. 47), and a miR482/Vv-sly-TAS5-like-3′D6 (+)-triggered LRR PHASI locus *VIT_12s0034g01350* (phase score 398.3; Supplementary Table 2b) were significantly up-regulated by high-fluence UV-B, consistent with observed trigger miRNA abundances (Fig. 2, Table 5). However, *TAS4c* (which has a 3′D4(-) variant that targets VviMYBA6/7 less well; mismatch at seed position 10; Supplementary Table 2b), despite evidence for being up-regulated by high fluence UV-B *per se* (like *TAS4ab;* Table 4), was expressed highly in both low fluence UV control and treatments compared to high fluence conditions (not shown), resulting in an overall negative coefficient for high-fluence effect like seen for miR403 family members (Tables 3,4). These results suggest an additional regulatory mechanism affecting *TAS4c* expression, which we speculate might involve the TAS4c-3′-D4(-) mismatches to target *MYBA6/A7* impacting a functional interaction with the hypothesized auto-regulatory loop (Tables 3,4).

**Table 6.**
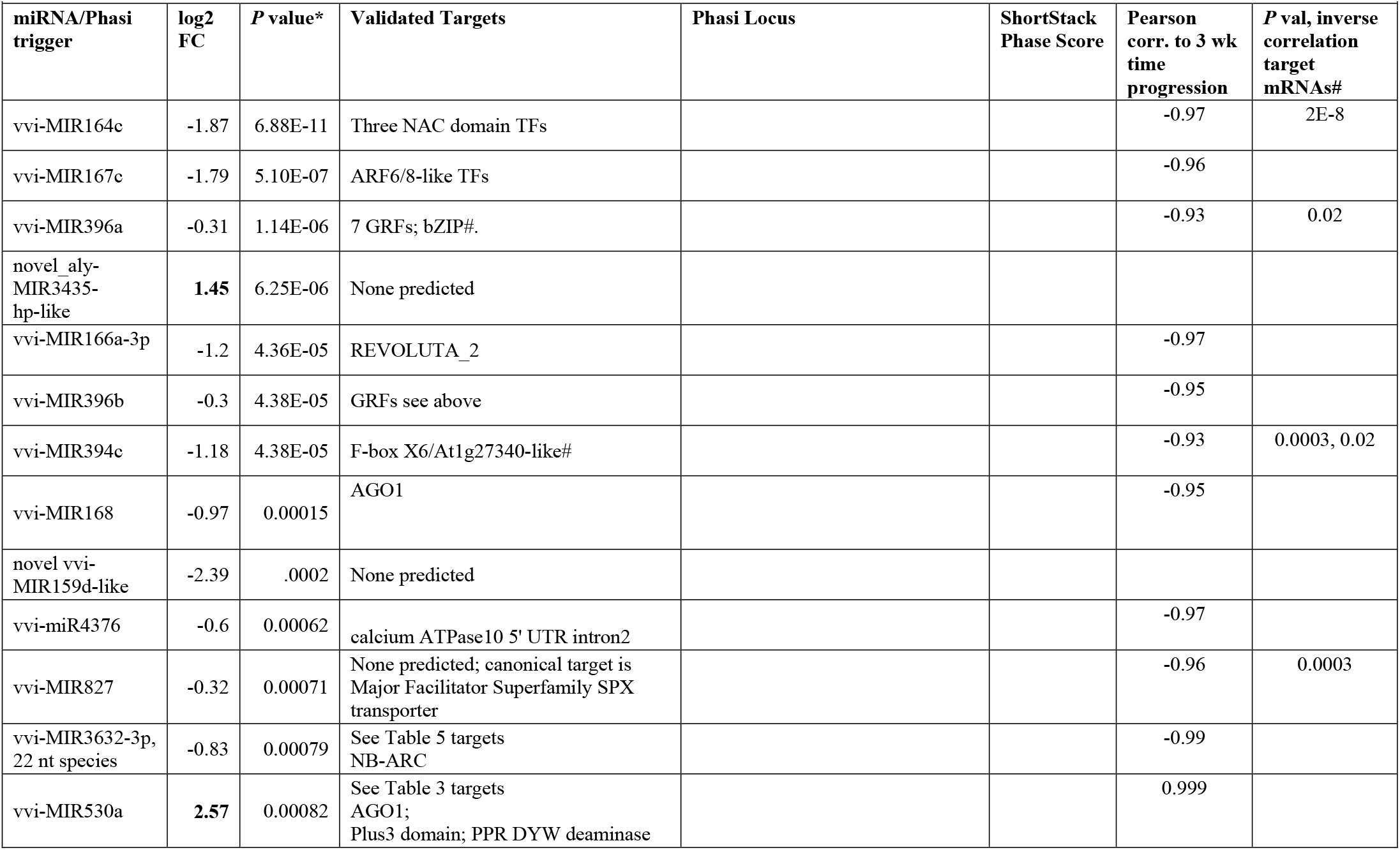
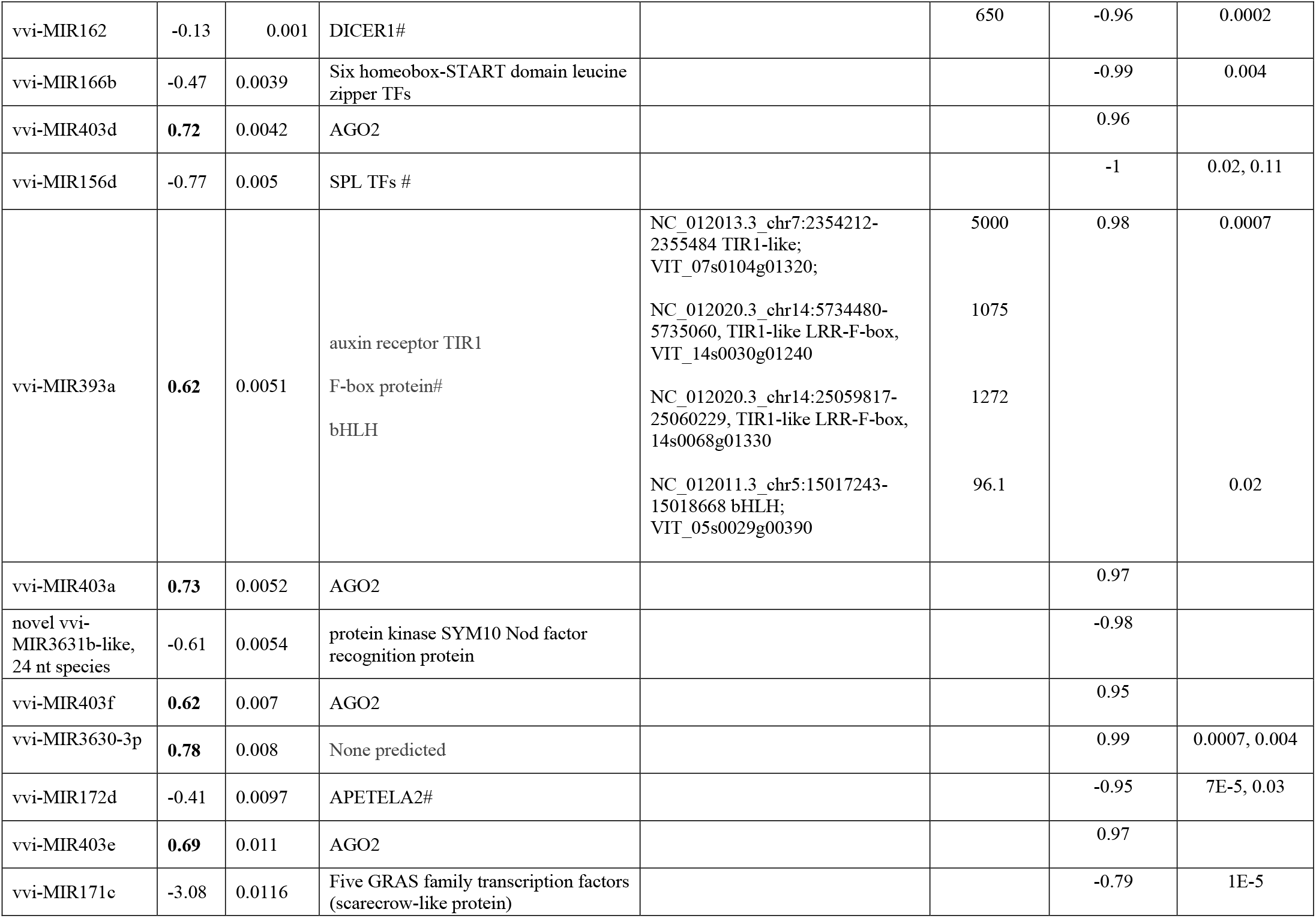
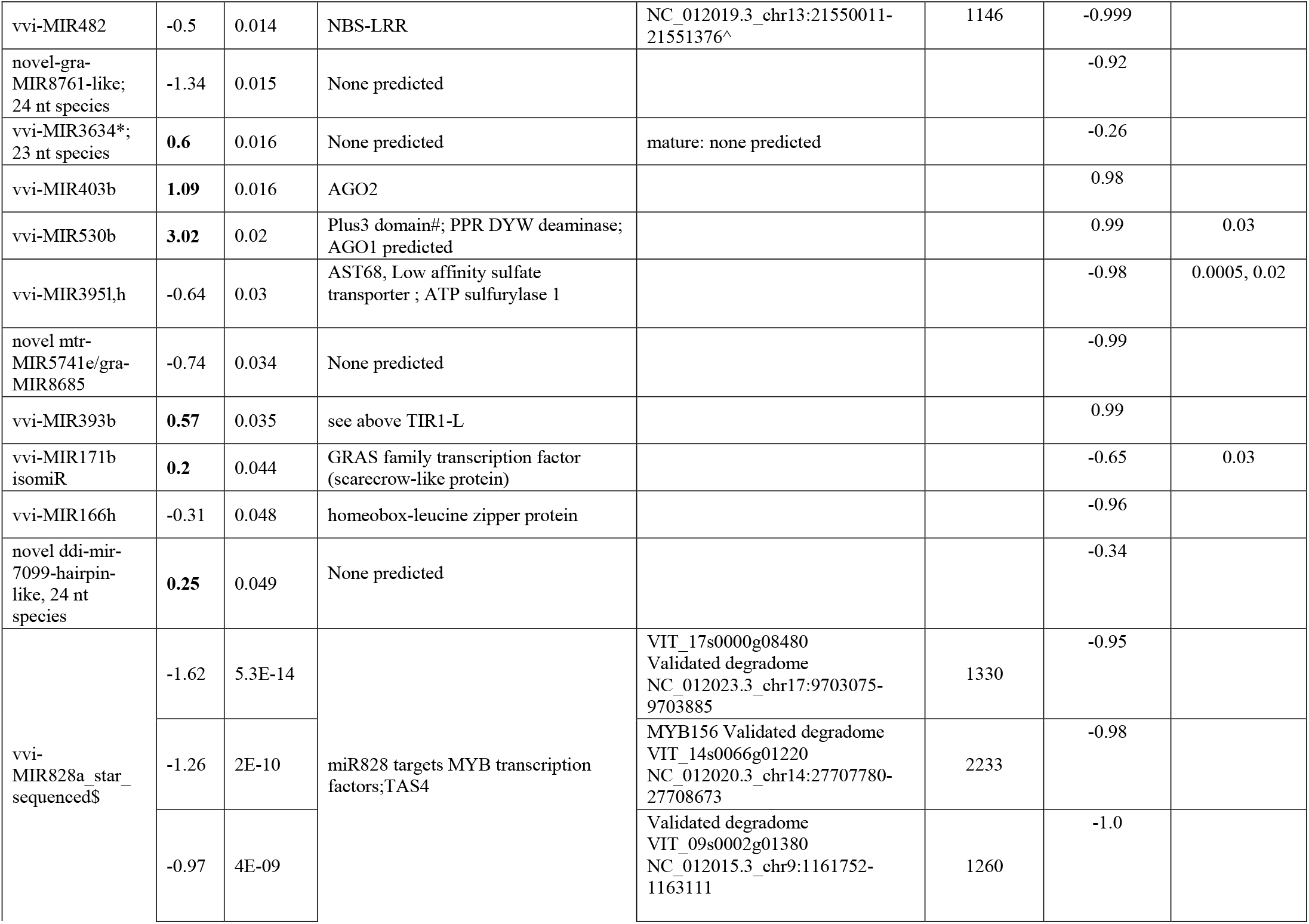
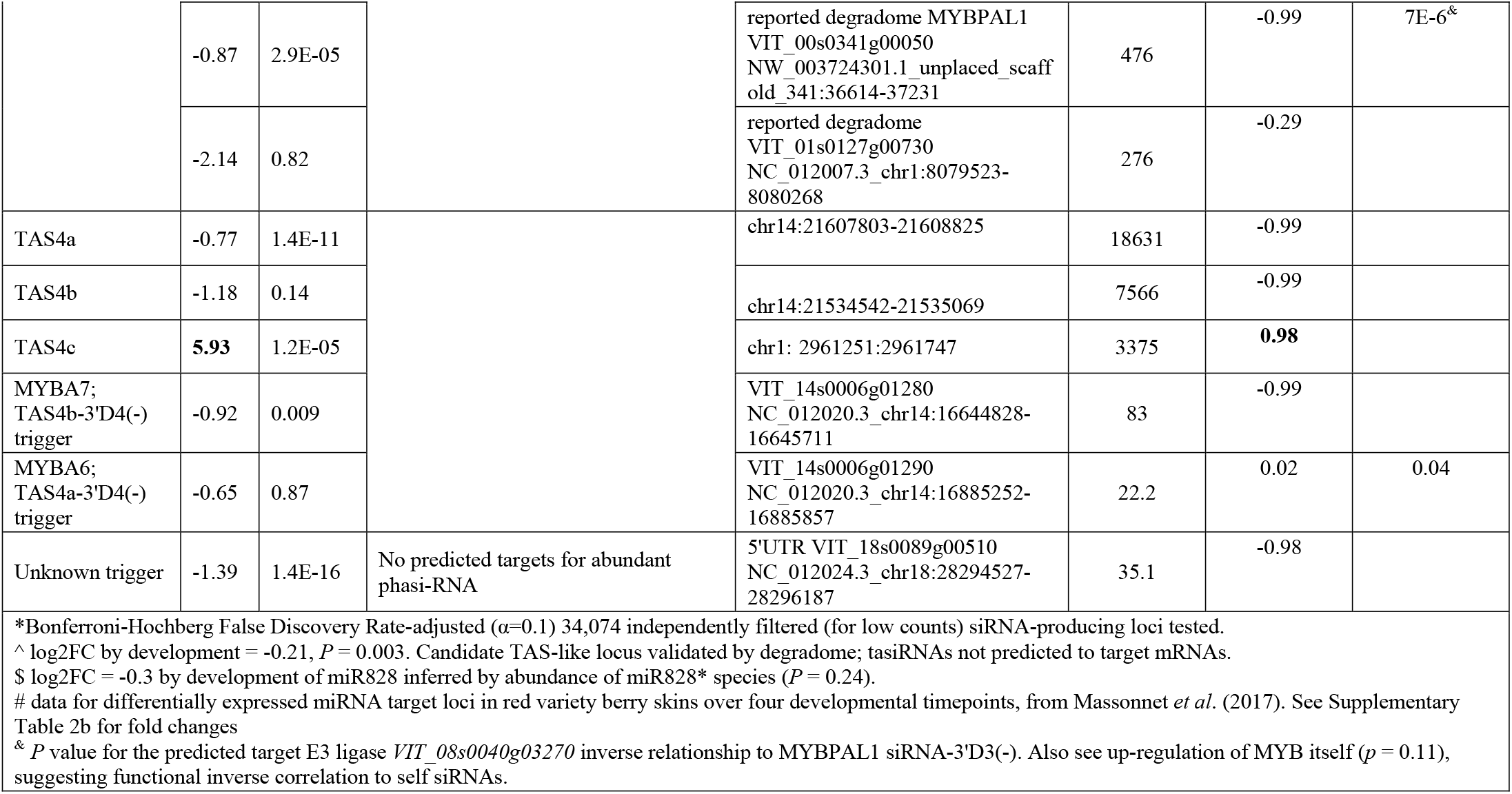
Differential expression of miRNAs in berries under different developmental stages. miRNAs that are up-regulated during berry development have log2FC column in **bold.**

### 3.4 Small RNAs differentially expressed during berry development

To characterize the overall accumulation of miRNAs and siRNAs affected by berry ripening, sRNA libraries constructed from berries (−3, 0, +3 and +6 WAV) harvested from greenhouse and fields were tested for differential expression across developmental time using the likelihood ratio test. By assuming (null model) there is no difference in miRNA expression profile between different berry development stages, we identified 48 miRNAs to be significantly differentially expressed at one or more developmental time points across environmental treatments (Table 6). As expected for phenotypic changes associated with veraison, there were observed profound dynamics in miRNA abundances. 21 miRNAs were up-regulated while 27 displayed a significant negative log fold-change during development. Linear regression (Pearson coefficient; Supplementary Table 3) as a metric to correlate time (three equally spaced time points before berry maturity at harvest) with normalized miRNA/siRNA abundances demonstrated a strong linear correlation across the developmental replicate time points (Table 6; Supplementary Table 3). This strong linear relationship held for nearly every *MIRNA* and PHASI locus even when fold changes were small and non-significant. This evidence demonstrates that the experimental parameters were highly reproducible and the results warrant inference of miRNA/siRNA functions towards validated targets as shown above. Co-expression analysis of our DE miRNAs (Table 6) with a published RNA-Seq experiment that reported significantly differentially expressed mRNAs at four comparable time points in five red-skinned varieties identified 15 cases of miRNA/target mRNAs with statistically significant (binomial test of chance) inverse correlations, supporting functional effects of miRNA changes during development (Table 6, Supplementary Table 2b).

*TAS4a, -b* and *-c* loci triggered by miR828, which in turn triggers phasing of *MYBA6, MYBA7*, and *MYBA5/MYB113-like* siRNA production via *TAS4ab*-3′-D4(-)(Supplementary File S4, pp. 38-43), showed differential expressions during berry development. *TAS4a* expression peaked at veraison when compared to -3 WAV, and decreased at +3 WAV and +6 WAV (Supplementary Table 3). *TAS4b* siRNAs did not display a pronounced up-regulation pattern across developmental stages. Interestingly, *TAS4c* reads increased at every developmental stage when compared to the previous stage (Supplementary Table 3), consistent with the observation (Table 5) *TAS4c* and miR403 were regulated differently than *TAS4ab* and most miRNAs, which decrease during berry development (Table 6). miR530ab may target 3′ UTR intronic regions of AGO1 orthologues, supported by GSTAr alignment data (Supplementary Table 2a) and miR393ab, which degradome evidence shows triggers phasi-RNA production and phasi-RNA auto- and *trans* cleavages (Si-Ammour *et al*., 2011) in *AUXIN TRANSPORT INHIBITOR RESPONSE1/TIR1* co-receptor mRNAs (Supplementary Tables 2a, 4; Supplementary Files S5, S4 p. 20), were also coordinately up-regulated during berry development like miR403 family members and *TAS4c* (Table 6).

## 4 Discussion

We have endeavored to characterize the miRNA and phasi-RNA space in response to UV-B light by target degradome validations, statistical inference of sRNA dynamics, and direct qRT-PCR measurement and/or meta-analysis of miRNA target mRNA abundances in published datasets from the same tissues. Shortstack ver3, by virtue of its multimapper functionality (Johnson *et al*., 2016), was our method of choice for quantifying locus-specific miRNA precursor (i.e. cluster) abundance as a proxy for mature miRNA and phasi-RNA abundances. This approach presumes that when a cluster is called as differentially expressed, the mature miRNA or most abundant phasi-RNA is the species changing in response to treatment or condition. Even if the subject sRNA ostensibly undergoes dynamic fluctuations in abundance, it cannot function as a negative effector of target gene activity without being loaded in a functional AGO-containing RISC, which along with base pairing to target mRNAs controls miRNA stability, activity, and subcellular retention (Li *et al*., 2013; Pitchiaya, Heinicke, Park, Cameron, & Walter, 2017; Speth, Willing, Rausch, Schneeberger, & Laubinger, 2013). It is remarkable that miR828, a 22 nt species with a very low estimated abundance (the sequenced star species = 2.6 rpm from 411 raw reads; thus miR828 being unsequenced ~ 0.01 rpm; Supplementary Table 3), manifested the highest *in vivo* slicing activities based on phasi-RNA abundances (~15,000 rpm) and phasing scores from *TAS4abc* and target *MYBs* (Supplementary Table 3; File S4, pp. 38-52) while on the other hand, we were unable to find slicing evidence for the relatively abundant miR397 (38 rpm), miR399 (42 rpm), and miR408 (a phasi-RNA producing locus with phase score of 393 and star species abundance of 883 rpm). It is speculated that modeling the unusually active (like miR828) and non-canonical effector:target duplexes containing bulges revealed by our degradome analyses (e.g. miR319, miR162, miR396, miR398, miR3627-iso; Supplementary Table 2a; File S4 pp. 4, 6, 24, 28-31, 57, respectively) programmed into a crystal structure of human AGO (Elkayam *et al*., 2012) may provide insights as the structures of plant and animal AGOs are conserved (Fátyol *et al*., 2016; Zhang *et al*., 2014).

Single nucleotide polymorphisms of sRNAs across mapping datasets could also be a confounding factor in interpreting our differential expression candidate miRNAs. Caution is warranted when interpreting phasi-RNA abundances (e.g. *TAS4a,b,c)* where there are many species generated, some of which can accumulate to high levels yet only one (e.g. *TAS4* 3′-D4[-]; Rock, 2013) is claimed as functional. The functional siRNA and miRNA may not be the major species in clusters or the one that changes significantly in response to stimulus, confounding interpretations. For *TAS4a*, the major species called by ShortStack is documented as 3′D1(+) (Supplementary Table 3, column “I”). Supplementary Table 3 and Supplementary File S4 (p. 38) provides direct evidence supporting the assumption that the claimed differentially expressed mature miRNAs and phasi-RNAs (e.g. *TAS4b,c* 3′D4[-]) are the major species changing their abundances *per se*. ShortStack output (Supplementary Table 3 columns Q and S) lists the “Major RNA” (i.e. most abundant) species and “Complexity”, calculated as (n distinct alignments)/(abundance of alignments). Low complexity values indicate loci dominated by just a few sRNAs. Although the ShortStack singular metrics as reported were the average across all libraries, the Major Species reported is invariably the mature miRNA species (or star, when it is claimed as functional as with miR3624-3p), and the Complexity reported is quite low, such that inference can be made that differentially expressed sRNAs are predictable. The average Complexity was 0.033 for claimed *MIRNA* clusters (data not shown), with several loci having zero complexity (see Supplementary File S6 for documentation of complexity vis-à-vis reads abundances for novel *MIRNAs)*. Complexity averaged 0.072 for *PHASI* loci across all claimed loci. For *TAS4a* where D1(+) was the major species the complexity was 0.002 and our 28 libraries accounted for ~11% of all mapped *TAS4a* reads (data not shown); for *TAS4b* the 3′-D4(-) siRNA was the major species and complexity was 0.001 wherein our libraries accounted for only 1% of all *TAS4b* reads. However *TAS4b* 3′D4(-) was highly abundant in the field samples and showed three to seven-fold increases in response to solar UV across all four developmental time points (Supplementary Table 3; Supplementary Fig. S2C, left panel). For *TAS4c*, 3′-D4(-) was also the major species and the measured complexity was 0.011; for this key hypothesized effector of *MYBA6* and *MYBA7* expression our libraries accounted for 91% of all *TAS4c* cluster reads. Taken together, these data support our view that statistical inference of miRNA and siRNA dynamics using clusters as proxy for *bona fide* effectors is reasonable and useful.

Initial prediction of UV-B-responsive miRNAs in Arabidopsis was based on computational analyses of UV-regulated co-expressed genes which shared promoter motifs with candidate *MIRNAs*, coupled with predicted target expression that inversely correlated with inferred miRNA expression (Zhou *et al*., 2007). UV-B responsive miRNAs were later identified in poplar (Jia *et al*., 2009), wheat (Wang *et al*., 2013), maize (Casati, 2013), and *Brassica rapa* for UV-A (Zhou, Fan, Li, Yan, & Xu, 2016). In maize, miR156 and miR529 are decreased and their several targets *Squamosa Promoter-Binding-like (SPL* transcripts) are increased after 8 hr UV-B in maize leaves (Casati, 2013). Wang *et al*. (2013) also reported down-regulation of miR156 by UV-B in wheat. We also observed inverse correlations between vvi-miR156 abundance and *SPL* target expressions in our experiments (Fig. 3) which were consistent with published datasets for UV-C and UV-B responses (Table 4, Supplementary Table 2b), but in contrast to maize and wheat we found vvi-miR156f and vvi-miR535c to be up-regulated in grapevine in response to UV-B (Table 3). It is possible that the low-fluence down-regulation effect on numerous miRNAs including miR156f (Table 4) could account for the differences between grape and maize or wheat, where a coordinated antagonist response pathway for low fluence UV-B may be involved. Previous work on grapevine leaves showed high fluence rate UV-B specifically modulates pathways and processes associated with oxidative and biotic stress, cell cycle progression, and protein degradation, whereas low fluence rate UV-B regulates the expression of responses to auxin, ABA, and cell walls (Pontin *et al*., 2010). Consistent with this notion is the differential UV-B fluence effects on *VviTAS3* phasi-RNAs (Table 5, Supplementary Tables 2b, 3, 4) and miR828 abundances (up in high-, down in low fluence; Fig. 2, Table 4), and with prior work showing *TAS4* is negatively regulated by ABA/sugar crosstalk signaling (Luo *et al*., 2012). The higher expression level and lack of UV-B induction of miR828 in *in* vitro-grown plantlets (Fig. 2A) in contra-distinction to berries (Fig. 2B-D) is consistent with a possible negative effect of increased ABA in leaves (Loveys, 1984).

### 4.1 A bimodal pathway operates under low-fluence UV-B regulating photomorphogenesis

UV-B signaling mechanisms affecting grape berry polyphenolics (flavor enhancers and fruit astringency components) have been characterized (Loyola *et al*., 2016; Martínez-Lüscher *et al*., 2013), but there is no description of grapevine UV-B responsive miRNAs and their significance for regulation of polyphenolic biosynthesis, stress responses, or berry development. We took a genome-wide systems approach to characterize expression of sRNAs from *in vitro-grown* plantlets, different developmental stages of berries (−3, 0, +3 and +6 WAV) irradiated with high/low fluence UV-B in green house, and different developmentally staged berries naturally irradiated (high fluence) in the field. Our analysis revealed a dynamic regulation of certain miRNAs following UV-B-induction and pervasive changes in miRNA profiles during fruit development. We have formulated a model (Fig. 4) to frame high fluence miRNA dynamic responses to UV-B in the context of post-transcriptional gene silencing processes impacting oxidative and nutrient (sulfur and phosphate, associated with anthocyanin accumulation) stress adaptation during grape berry development. We suggest that these high fluence UV-B response processes may be different between monocots and dicots, like the case for *MIR828/TAS4* regulation of anthocyanin biosynthesis (Rock, 2013) based on three grounds: 1) observed differences between our results for deeply conserved and functionally significant miR156 changes (Fig. 3, Table 4) in contradistinction of results in maize and wheat (Casati, 2013; Wang *et al*., 2013). 2) The proposed coordinate antagonist response pathway for UV-regulated miRNAs revealed by the low fluence UV-B experiment (Fig. 2, Table 4) affecting predominantly dicot-specific miRNAs (Tables 2, 3, 5). 3) Differences in overrepresented gene ontology categories between eudicots and monocots suggest the evolution of oxygen and radical detoxification (i.e. UV associated) and RNA silencing processes mark key divergence points between dicots and monocots (Lee *et al*., 2011). Our results (Table 4) suggesting a bimodal/inverse pathway for miRNA dynamics with no overlap between the miRNAs regulating photomorphogenic and defense pathways in response to low fluence UV-B are in accordance with those of Yoon *et al*. (2016) and Pontin *et al*. (2010), who showed low levels of UV-B irradiation result in photomorphogenic and auxin responses rather than induction of defense- and abiotic oxidative stress-related genes.

**Figure 4.**
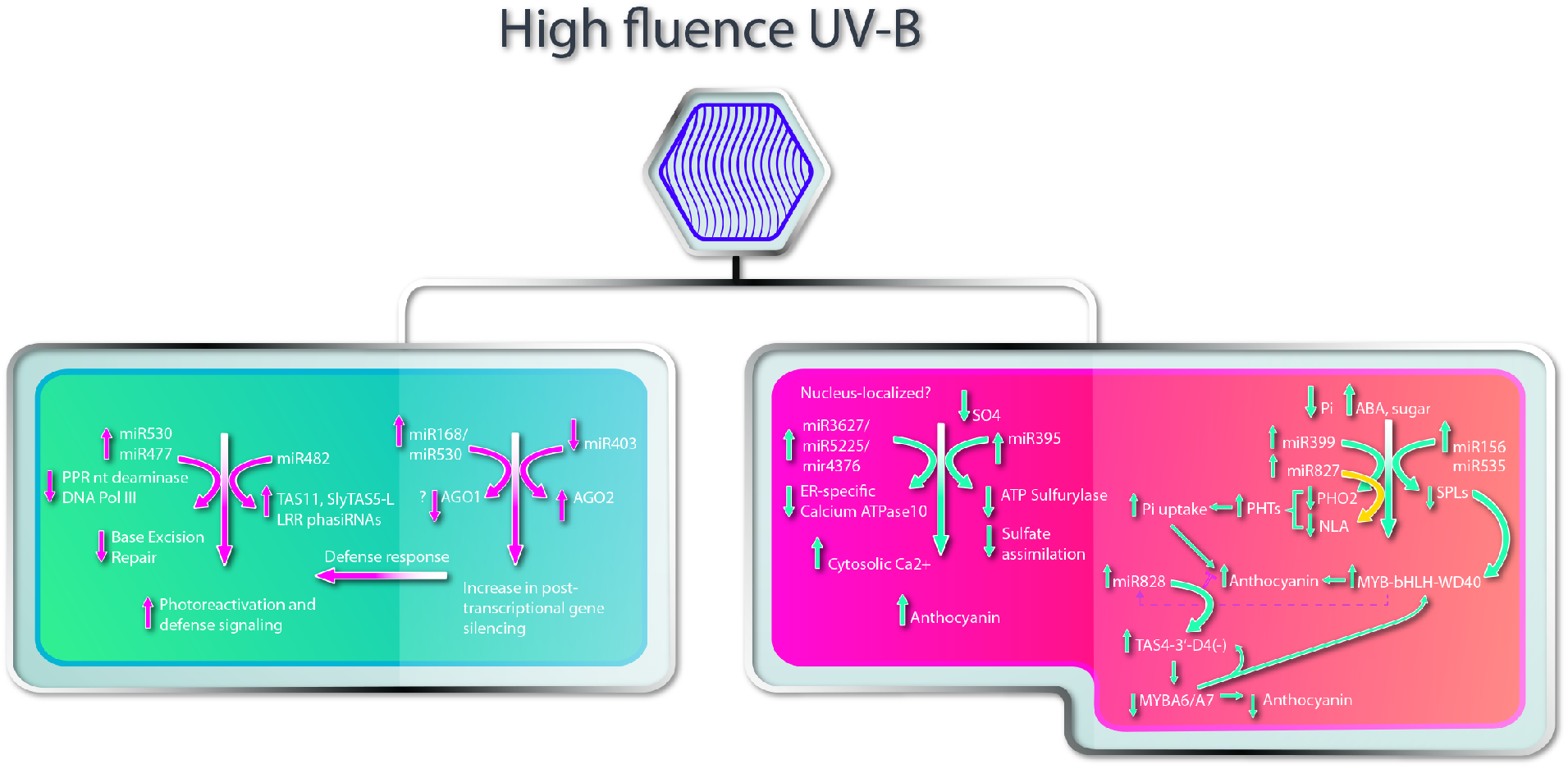
Model of UV-B effects on grape berry stress responses and development regulated by miRNAs. Defense, DNA repair and accumulation of anthocyanin are the major responses regulated by UV-B responsive miRNAs. Increase in miR168/miR530 facilitate post-transcriptional gene silencing activity. The increase in PTGS activity would in turn facilitate targeting of *LRR* genes by miR482 and regulate the defense genes. miR477 family members target *DNA Pol III* involved in base excision repair (BER) and facilitates photo-reactivation and nucleotide excision repair (NER) to remove UV-B induced DNA lesions. Upregulation of miR3627/4376/5225 by UV-B targets ER-specific *Calcium ATPase10* and facilitate increase in cytosolic Ca^2+^ levels and thereby increased anthocyanin accumulation. miRNAs shown to be regulated by sulfur (miR395) and phosphorous starvation (miR399; miR827) are also responsive to UV-B radiation and facilitate anthocyanin accumulation. miR156/miR535 target *SPL* genes and increase the accumulation of MYB-bHLH-WD40 TFs and anthocyanin. Increase in MYB-bHLH-WD40 TFs would trigger the conserved auto-regulatory loop involving miR828/TAS4 to regulate *MYBA6/A7* levels and thereby anthocyanin levels.

### 4.2 Polyphenolic pathway and cellular transcription are targets of UV-B responsive miRNA

The evolutionarily related miR5225, miR3627, and miR4376 (offset by 4 nt) were up-regulated upon UV-B irradiation (Tables 2, 3) and target 5′ UTR intron2 region of *Calcium-transporting ATPase10* pre-mRNA (Supplementary Table 2a; Supplementary File S4, pp. 56-57) in the nucleus to modulate free cytosolic Ca^2+^ level and *CHS* expression (Fig. 4). Frohnmeyer, Loyall, Blatt, and Grabov (1999) demonstrated that UV-B induces increases in free cytosolic Ca^2+^ levels associated with UV-B induction of *CHS* in parsley cells, and Loyola *et al*. (2016) demonstrated UV-B induction of *CHS3* in irradiated grapevine plantlets. Calcium transporting ATPase10 is known to be involved in transporting free cytosolic Ca^2+^ to the endoplasmic reticulum (Geisler, Frangne, Gomes, Martinoia, & Palmgren, 2000). Considering nutrient effects on a miRNA regulatory network impacting anthocyanin metabolism in response to stresses, vvi-MIR399i was identified to be up-regulated by high-fluence UV-B; its target *PHO2* encodes a ubiquitin-conjugating E2 enzyme that upon repression by miR399 (Supplementary Table 2a) in response to UV would promote increases in the stability of phosphate transporter and P_i_ uptake (Lin *et al*., 2008) (Fig. 4). Consistent with the model, phosphate deficiency increases anthocyanin content in grapevine cell culture (Yamakawa, Kato, Ishida, Kodama, & Minoda, 1983). vvi-MIR395 is a 5′ RLM-RACE-validated slicer of *ATP sulfurylase 1/GSVIVT01018057001/VIT_05s0020g04210* supported by GSTAr alignment prediction (Supplementary Table 2a), which showed a strong and significant inverse correlation between miRNA and target mRNA abundances in response to cold stress in grapevine (Sun *et al*., 2015). ATP sulfurylase along with validated target *VIT_18s0001g04890/AST68* low affinity sulfate transporter (Supplementary Table 2a, File S4 p. 22) function to increase sulfur transfer to aerial parts under sulfur starvation (Kawashima *et al*., 2011). MIR395 up-regulation in response to drought stress in rice (Zhou *et al*., 2010) and salinity stress in *Zea mays* (Ding *et al*., 2009) is consistent with our evidence for a role in abiotic stress responses including UV-B. Sulfur starvation increases anthocyanin levels in rice (Lunde *et al*., 2008) and in grapevine sulfur-dependent oxidative response effectors GLUTATHIONE S TRANSFERASE and glutaredoxin are induced during veraison (Terrier *et al*., 2005). Also, different GSTs have been recently related to flavonoid transport (Pérez-Díaz, Madrid-Espinoza, Salinas-Cornejo, González-Villanueva, & Ruiz-Lara, 2016), with VviGST1/4 having major roles in anthocyanin transport (Conn, Curtin, Bézier, Franco, & Zhang, 2008; Gomez *et al*., 2011). UV-B induces the accumulation of vvi-MIR535c and vvi-MIR156f, the latter of which we show (Fig. 3; Supplementary Table 2a; File S4 p. 1) and impute (Table 4) to coordinately down regulate *SPL* transcripts. Up-regulation of miR535 (Table 3) also may contribute to observed down-regulation of *SPLs*, although a bulge at nt 6 of the seed region and lower degradome p-values qualify this assertion (Supplementary Table 2a; 2b). Notwithstanding, there were compelling degradome evidences for slicing by several miRNAs of targets with bulges in the seed region: for miR396 canonical target *GRF*s, miR162 target *DICER1*, miR477 novel targets, and four novel copper-associated miR398 targets with large bulges, analogous but distinct and more extreme than the 6 nt mRNA bulge described for Arabidopsis *Blue Copper Binding plastocyanin-like* (Brousse *et al*., 2014) (Supplemental Table 2a, File S4 pp. 6, 24, 28-33, 35). Several groups have reported *SPL* transcripts in grapevine targeted by both miR156 and miR535 (Carra *et al*. 2009; Han *et al*., 2016; Pantaleo *et al*. 2010). *SPL* transcript reduction increases anthocyanin levels in Arabidopsis (Gou *et al*., 2011) and our functional co-expression analysis supports that UV-B regulation of miR156 is an important component of a regulatory network for anti-oxidant and UV-protective polyphenolic biosynthesis (Fig. 4).

vvi-MIR530 can possibly target PPR DYW deaminase *VIT_18s0001g13940* as shown by GSTAr alignment (Supplementary Table 2a), involved in chloroplast RNA editing (Wagoner, Sun, Lin, & Hanson, 2014) or mitochondrial dysfunction associated with oxidative stress that generates highly mutagenic 5-hydroxymethyluracil and 5-hydroxyuracil lesions (Kow, 2002). The preponderance of transcription factor and TAS/PHASI targets of UV-B regulated miRNAs in grapevine and across species supports a model (Fig. 4) integrating nodes of conserved regulatory networks important for integrating abiotic stress and developmental programs.

### 4.3 Polyphenolic pathway and defense genes are targets of UV-B responsive tasiRNAs

Recent papers report the role of miR858 in immunity of Arabidopsis to infections by nematodes (Piya *et al*., 2017) and fungal pathogens by negative regulation of flavonoid-specific *MYB* TF genes (Camargo-Ramírez, Val-Torregrosa, & San Segundo, 2018). Other researchers have invoked the ‘two-hit’ model (Axtell *et al*., 2006) to explain observed phasi-RNA production of miR828- and miR858-dual targeted *MYBs* involved in proanthocyanidin and anthocyanin biosynthesis, based on weak predicted binding affinity of miR858 in a conserved domain 12 nt upstream of miR828 binding sites (Guan *et al*., 2014; Xia, Zhu, An, Beers, & Liu, 2012; Zhu *et al*., 2012). Several grapevine *MYBs* with a conserved miR858-miR828 dual binding domain as described in peach, apple, cotton, and Arabidopsis (Guan *et al*., 2014; Xia *et al*., 2012; Zhu *et al*. 2012) showed compelling evidence of slicing by miR828 that spawns phasi-RNAs in register, however these targets only showed marginal evidence of miR858 slicing that correlates with high (4.5-6.5) Allen scores (Supplementary File S4, pp. 44, 47, 49, 50). Although it is plausible that dual auto-regulatory loops for *miR828:MYBA6/7* (Rock, 2013) and miR858:MYB82 (Piya et al., 2017) operate coordinately in grapevine to regulate anthocyanin and flavonol biosynthesis, respectively like in Arabidopsis, our functional analysis supports miR828 being the dominant effector module. It is noteworthy that for *MYBA6/7/A5-MYB113-like* slicing by 21 nt TAS4ab-3′D4(-) and production of abundant phased registers downstream, there is no obvious trigger mechanism as has been described for the ‘two-hit’ and 22_1_ hit miRNA models for phasi-RNA production (Axtell et al., 2006). It will be interesting to determine if *MYBA5/MYB113-like* is targeted by both miR828 and TAS4-3′D4(-) as in Arabidopsis (Rajagopalan et al., 2006; Luo et al., 2012).

vvi-MIR828-triggered *TAS4abc* 3′D4(-) tasiRNAs (Supplementary Table 2a; Supplementary File S4, pp. 38-40) and primary *TAS4* transcripts are up-regulated by high fluence UV-B in greenhouse and field (Table 4, Fig. 3E, Supplementary Fig. 2B, C, D) which is in line with the function of a conserved auto-regulatory feedback loop proposed by Hsieh *et al*. (2009) and Luo *et al*. (2012). Increases in VvMYBA6/7 induce *TAS4* (and *MIR828)* transcription and thereby trigger a feedback repression of *MYBs* targeted by miR828 and *TAS4*. Novel *TAS11* and vvi-slyTAS5-like loci are triggered by miR482 up-regulation under high fluence UV which generate several phasi-RNAs from multiple registers that target numerous *LRR* genes which themselves undergo UV-B dependent siRNA amplification (Tables 2,3, 5) and subsequent functional silencing (Table 4, Fig. 3D; Supplementary Table 2b; Supplementary File S4, p. 37). The repression of defense genes by miRNAs until the plant is exposed to severe stress is an adaptive mechanism hypothesized to have evolved to keep the defense response under post-transcriptional control to conserve energy by preventing LRR protein synthesis (Shivaprasad *et al*., 2012). A domain of the nuclear Armadillo-type found in Cap Binding large subunit and LRR variants is targeted by D6(+) phase of vvi-slyTAS5-like based on degradome evidence (Supplementary Table 4). Mutations in *cap binding protein20* and *cbp80* are ABA hypersensitive, drought tolerant, and have abnormal miRNA biogenesis (Gregory *et al*., 2008; Papp, Mur, Dalmadi, Dulai, & Koncz, 2004). It is plausible that targeting of a cap binding complex by elevated miR482/*TAS* phasi-RNA expression may be an adaptive mechanism for UV-B response/tolerance and a crosstalk mechanism linking UV-B fluence dynamics of miRNAs (Table 4) to ABA antagonism of *MIR828/TAS4* feedback homeostasis (Luo *et al*., 2012).

### 4.4 Oxidative stress by high fluence UV-B: combated by miRNAs?

The 18 differentially expressed miRNAs identified (Table 5) as changing under high versus low fluence *per se* (including correlated changes in greenhouse controls) were more than found when UV-B radiation across all environments including the field was tested as a single parameter (Table 3). This is possibly because of the noise associated with different (i.e. field) environments. However, Pontin *et al*. (2010) observed the same degree of increased effects of high-versus low fluence UV-B changes in a grapevine leaf transcriptome comparison. One upshot from the alternative perspectives afforded by the controlled greenhouse experiments was the finding that certain miRNAs/siRNAs (e.g. miR403, miR530, *TAS4c;* Tables 5, 6) are regulated differently and possibly coordinately by UV and development. Six miRNAs (five of them novel; Supplementary File S6) belonging to the *MIR477* family were up-regulated by UV-B and had star species which were much more abundant. vvi-miR477b is predicted to uniquely target *RepC/DNA Pol III epsilon subunit τ* associated with nucleotide excision/mismatch repair pathways. Interestingly, *MIR477, MIR482*, and *MIR3627* are conserved across woody fruit trees (Solofoharivelo, van der Walt, Stephan, Burger, & Murray, 2014) including grapevine. We speculate the prevalence of vvi-MIR477 family members induced by high fluence UV-B may indicate a regulatory role in countering oxidative stress. UV-B radiation is known to induce cyclobutane pyrimidine dimers that kill cells by blocking DNA replication (Ries *et al*., 2000). Photorepair by photolyase and nucleotide excision repair (NER) are the major pathways in plants for repairing UV-B-induced DNA damage (Tuteja, Singh, Misra, Bhalla, & Tuteja, 2001), while base excision repair (BER) involving RepC/DNAPol III epsilon (predicted target of miR477b) is involved in repairing oxidized or hydrated bases and single-stranded breaks caused by ionizing radiation (Gill, Anjum, Gill, Jha, & Tuteja, 2015). We speculate that the observed up-regulation of *vvi-MIR477* family members may be an adaptation to reduce BER mechanisms and facilitate photoreactivation and NER to remove UV-B induced DNA lesions (Fig. 4).

Liang, Li, He, Wang, and Yu (2013) demonstrated that in addition to miR168, AGO1 is a target of miR530 in papaya and a predicted target of miR530 in *Salvia miltiorrhiza* (Shao & Lu, 2013). We predict that two grape AGO1 orthologues could also be targets of vvi-miR530-isomiR (Supplementary Table 2a). Harvey *et al*. (2011) demonstrated AGO2 acts as a second layer of control in antiviral defense which is activated when AGO1 is suppressed. When AGO1 is active it represses *AGO2* via miR403 activity. Interestingly, AGO2 induction by *Pseudomonas syringae* pv. *tomato* bacterial infection without suppression by its negative effector miR403 was observed in the *ago1-27* mutant, suggesting a post-transcriptional regulation of *AGO2* by bacteria which is mechanistically different from the induction triggered by viruses (Zhang *et al*., 2011). Our assay of *AGO2* transcript abundances in samples from low- and high fluence UV-B greenhouse-treated berries showed inverse trends to documented miR403 dynamics (Table 4) but were not statistically significant (data not shown). Based on the observed coordinate down-regulation of all vvi-miR403 family members and imputed up-regulation (albeit not statistically significant; *p* = 0.17) of validated target *AGO2* in response to UV-B (Tables 4, 5; Supplementary File S4, p. 34), we speculate on a third component to the model that might operate in UV-B-radiated grape in which AGO2 could play a role in combating UV oxidative stress without inactivation of AGO1 (Fig. 4).

### 4.5 Auto-regulatory loop involving miR828-7¾Ŵ-MYBA6/A7/A5-MYB113-like is conserved in berry development

Forty-eight miRNAs were identified as differentially expressed across berry development (Table 6). Our finding that miR396, miR172, and miR171 are down -regulated during berry development (Table 6) and correlated with meta-analysis showing up-regulation of some targets (Supplementary Table 2b; data of Massonet *et al*., 2017) is consistent with the results of Mica *et al*. (2010) who quantified miRNAs during berry development using a custom microarray. Our specific rationale for studying miRNAs/phasi-RNAs is their involvement in polyphenolic pigmentation, one of the most important traits for grape berry ripening and subject to nutrient and phytohormone effects (Vanhaelewyn, Prinsen, Van Der Straeten, & Vandenbussche, 2016). UV-B radiation reduces berry weight and increases anthocyanin content (Berli *et al*., 2010; Berli, Fanzone, Piccoli, & Bottini, 2011). In addition to phosphate deficiency (Yamakawa *et al*., 1983), increases in ABA and response effectors increase anthocyanin levels (Hiratsuka *et al*., 2001; Koyama, Sadamatsu, & Goto-Yamamoto, 2010; Luo *et al*., 2012). Interestingly, UV-B radiation does not affect berry ABA content whereas leaves exposed to UV-B increase ABA levels (Berli *et al*., 2010; Berli & Ruben, 2013). The vegetative anthocyanin loci *MYBA6,MYBA7*, and presumably *MYBA5/MYB113-like* and their regulator *HY5* are up-regulated at early stages when exposed to UV-B and despite their expression declines with berry ripening, this behavior is delayed in response to UV (Supplementary Fig. S2E, F; Loyola *et al*. 2016; Matus *et al*., 2017). Pilati *et al*. (2017) reported up-regulation of *MYBA7* when berries are treated with ABA for 20 h but not when the treatment was prolonged to 44 h. This observation is consistent with the notion that *MYBA7* is induced directly by ABA but does not have a role in berry ripening *per se. vvi-MIR828* and *TAS4* are validated here to regulate the *MYBA6, MYBA7*, and *MYB5/MYB113-like* levels. The auto-regulatory feedback loop is shown to function in grapevine similar to Arabidopsis (Hsieh *et al*., 2009; Luo *et al*., 2012), whereby developmental induction early in berry development of MYBA6 and MYBA7 correlates with observed elevated miR828 and high-fluence UV-B inductions at the green berry and veraison stages (Fig. 2B, D) with concordant temporal accumulations of *TAS4abc* primary transcripts and derivative siRNAs (Fig. 3E, Supplementary Fig. S2, Supplementary Fig. S3, Supplementary File S4, p. 38-43).

Interestingly, *TAS4c* which has two single nucleotide polymorphisms compared to the TAS4c-3′D4(-) effector species reported earlier (Rock, 2013) displayed a trend of increasing reads abundance with advancement of berry ripening (Supplementary Table 3). This result is intriguing because earlier studies on *Vitis amurensis* and *Vitis vinifera* (cv. Pinot Noir) grape datasets could not detect appreciable *TAS4c* phasi-RNA expression (Rock, 2013). It will be critical to ascertain if the trait of *TAS4c*-3′D4(-) expression late in berry development (Table 6) is unique to cv. Cabernet Sauvignon. *TAS4ab*-3′D4(-)-triggered *MYBA6* and *MYBA7* phasi-RNAs corresponding to the sixth phase are the most abundant (Rock, 2013). Similar to earlier work we find here that the sixth phase of *MYBA6/7* phasi-RNAs are the major species (Supplementary Table 3) where *MYBA7* phasi-RNA D6(+) is significantly down-regulated during berry development, in stark contrast to *TAS4c3′* -D4(-) significant up-regulation (Table 6). Further evidence consistent with the functioning of this feedback loop comes from RNA-Seq transcriptome data for several red-skinned berries (Massonnet *et al*., 2017). The reported significant up-regulation of *MYBA6* mRNA (Supplementary Table 2b) at pre-veraison to veraison transition is *inversely* correlated with observed down regulation of *TAS4ab* and downstream *MYBA6/7* phasi-RNAs (Table 6), suggesting a positive feed-forward activity for the TAS4ab 3′D4(-) tasi-RNAs slicing *MYBA6/7* and *MYBA5/MYB113-like* (Supplementary Table 2a; Supplementary File S4). The subsequent wholesale inversion of the *MYBA6* target mRNA: *TAS4ab* primary transcript and tasi-RNA relationships (i.e. both significantly down; Supplementary Table 2b, Supplementary Fig. S2, Supplementary Fig. S3) at berry maturity is parsimonious with the predicted silencing functions of TAS4ab-3′D4(-) on *MYBA6/7* late in berry development. Matus *et al*. (2017) showed high level expression and significant UV-B induction of *MYBA6/7* before veraison and complete down-regulation of *MYBA7* (but not *MYBA6*, which was delayed) during and after veraison. Our results of differential UV-B fluence effects on miR828 (Fig. 2B-D) and developmental stage-specific *TAS4a-c* expressions (Tables 4, 6) suggest that the auto-regulatory circuit may function to fine-tune expression of *MYBA6/7* in response to UV-B and during berry skin maturation. Our results reveal complex activities of *MIR828/TAS4* in response to environmental and developmental queues to post-transcriptionally silence target effectors, and underscore the importance of post-transcriptional controls in the observed delays by UV-B exposure on repression of *MYBA6* late in berry development. These exploratory results and meta-analysis provide a basis for systems approaches to better understand non-coding RNA functions in response to UV light and strategies for improving winegrape production of anti-oxidant flavor enhancers.

## Conflict of interest

The authors declare that the research was conducted in the absence of any commercial or financial relationships that could be construed as a potential conflict of interest. A U.S. patent application was filed by Texas Tech University System (C.D. Rock & Q. Luo. “siRNAs compositions and method for manipulating berry ripening”, USPTO Publication No. US-2014-0075596-A1).

## Author Contributions

conception and design of the work: JTM, PAJ, JAA, CR; acquisition, analysis, interpretation of data: SS, CR, RL, JTM; wrote the manuscript: SS, CR; revised the manuscript: JTM, RL.

## Funding

FAPESP-SPRINT Brazilian-TTU Joint Program and TTU-VPR Open Access Publication Initiative (to CR), Comisión Nacional de Investigación Científica y Tecnológica, Chile (CONICYT; Ph.D. grant no. 21120255 to RL), Fondo Nacional de Desarrollo Científico y Tecnológico (FONDECYT postdoctoral grant no. 3150578 to RL and 1150220 to PAJ), NC130030 “Millennium Nucleus Center For Plant Systems And Synthetic Biology” and NM-BSBSV (to PAJ).

## Acknowledgements

The authors thank Glenn Hicks and John Weger at the Institute for Integrative Genome Biology UC Riverside for Illumina Nextseq500 sequencing, Rao Kottapalli at the Texas Tech Center for Biotechnology and Genomics for MiSeq sequencing, and the TTU High Performance Computer Center for support in use of the Quanah supercluster.

---

**Supplementary Figure S1.** Collective size distribution of adapter-trimmed reads in the 28 UV experimental sRNA libraries.

**Supplementary Figure S2.** Functional validation of differentially expressed miRNA/tasiRNA activities under high fluence solar UV-B in the field by qRT-PCR of target genes. **A)** Schematic alignment of *vvi-TAS4a, -TAS4b* and *-TAS4c* primary transcript sequences flanking the miR828 binding and *TAS4* 3′ D4(-) phasi-RNA positions, and location of locus-specific primers used for qRT-PCR. **Panels B-D:** Expression profiles of *TAS4a, TAS4b* and *TAS4c* siRNAs (left panels) and their cognate primary undiced transcript abundances. **B)** *TAS4a* 3′ D1(+) *trans* acting siRNA expression profile and the expression profile of *TAS4a* primary transcript. **C)** *TAS4b* 3′ D4(-) *trans* acting siRNA expression profile and the expression profile of *TAS4b* primary transcript. **D)** *TAS4c* 3′ D4(-) *trans* acting siRNA expression profile and the expression profile of *TAS4c* primary transcript. **E)** TAS4-3′D4(-) target *MYBA6* phasi-RNA-3′D6(+) expression profile and the expression profile of *MYBA6* mRNA. **F)** TAS4-3′D4(-) target *MYBA7* phasi-RNA-3′D6(+) expression profile and the expression profile of *MYBA7* mRNA. Error bars are s.d. Asterisks (*) denote significant differences based on analysis of variance (n=4 biological replicates) and comparisons using the Tukey-Kramer honestly significance test (HSD; *p* < 0.05). **Insets, B-F:** Target:miRNA/tasi-RNA binding positions (mRNA nucleotide coordinates) and primer binding sites (arrows) are depicted to scale. Target:miRNA binding depicted in Red; Target:tasiRNA binding depicted in Green; Target:phasiRNA binding depicted in Pink.

**Supplementary Figure S3.** miR403f mature expression profile and the expression profile of *MIR403f* primary transcript, showing concordant inductions during berry development. Error bars are s.d. Asterisks (*) denote significant differences based on analysis of variance (n=4 biological replicates) and comparisons using the Tukey-Kramer honestly significance test (HSD; *p* < 0.05). **Inset:** *pre-MIR403f*:miRNA binding position (mRNA nucleotide coordinates) and primer binding sites (arrows) are depicted to scale. Target:miRNA binding depicted in Red.

**Supplementary Table 1a and 1b.** small RNA library (a) and degradome (b) quality control parameters.

**Supplementary Table 2a and 2b. a)** List of validated miRNA targets by CleaveLand4.4.**b)** Metaanalysis of validated miRNA targets for three published datasets for transcriptome changes of berry skins in response to: UV-C (Suzuki *et al*., 2015), UV-B (Carbonell-Bejerano *et al*., 2017) and development (Massonnet *et al*., 2017).

**Supplementary Table 3.** DESeq2 output and associated ShortStack raw Counts dataset parameters for rows in Tables 2,3,5,6 and normalized reads per 20M for visualization of compared effects.

Supplementary Table 4. PhaseTank output of degradome *TAS* loci targets and miRNA/phasi-RNA trigger predictions.

**Supplementary File S4.** CleaveLand-validated miRNA target T plots.

**Supplementary File S5.** PhaseTank Align and Cascades Output, UV-regulated *PHASI* loci.

**Supplementary File S6.** Novel *MIRNA* loci characterized de novo by ShortStack, drawn from reads of libraries constructed from UV-B treated samples.

**Supplementary File S7.** List of primers and parameters used in this study.

## References

1. Addo-Quaye C., Miller W. & Axtell M.J. (2009) CleaveLand: a pipeline for using degradome data to find cleaved small RNA targets. Bioinformatics 25, 130–131. doi: 10.1093/bioinformatics/btn604

2. Alabi O.J., Zheng Y., Jagadeeswaran G., Sunkar R. & Naidu R.A. (2012) High-throughput sequence analysis of small RNAs in grapevine (Vitis vinifera L.) affected by grapevine leafroll disease. Molecular Plant Pathology 13, 1060–1076. doi: 10.1111/j.1364-3703.2012.00815.x.

3. Axtell M.J., Jan C., Rajagopalan R. & Bartel D.P. (2006) A two-hit trigger for siRNA biogenesis in plants. Cell 127, 565–577.

4. Belli Kullan J., Lopes Paim-Pinto D., Bertolini E., Fasoli M., Zenoni S., Tornielli G.B.,…,Mica E. (2015) miRVine: a microRNA expression atlas of grapevine based on small RNA sequencing. BMC Genomics 16, 393.

5. Berli F.J., Fanzone M., Piccoli P. & Bottini R. (2011) Solar UV-B and ABA are involved in phenol metabolism of Vitis vinifera L. increasing biosynthesis of berry skin polyphenols. Journal of Agricultural and Food Chemistry 59, 4874–4884.

6. Berli F.J., Moreno D., Piccoli P., Hespanhol-Viana L., Silva M.F., Bressan-Smith R.,…,Bottini R. (2010) Abscisic acid is involved in the response of grape (Vitis vinifera L.) cv. Malbec leaf tissues to ultraviolet-B radiation by enhancing ultraviolet-absorbing compounds, antioxidant enzymes and membrane sterols. Plant, Cell and Environment 33, 1–10.

7. Berli F.J. & Ruben B. (2013) UV-B and abscisic acid effects on grape berry maturation and quality. Journal of Berry Research 3, 1–14

8. Brosché M., Schuler M.A., Kalbina I., Connor L. & Strid A. (2002) Gene regulation by low level UV-B radiation: identification by DNA array analysis. Photochemical & Photobiological Sciences 1, 656–664.

9. Brousse C., Liu Q., Beauclair L., Deremetz A., Axtell M.J. & Bouché N. (2014) A non-canonical plant microRNA target site. Nucleic Acids Research 42, 5270–5279.

10. Borevitz J.O., Xia Y., Blount J., Dixon R.A. & Lamb C. (2000) Activation tagging identifies a conserved MYB regulator of phenylpropanoid biosynthesis. The Plant Cell 12, 2383–2394.

11. Camargo-Ramírez R., Val-Torregrosa B. & San Segundo B. (2018) miR858-mediated regulation of flavonoid-specific MYB transcription factor genes controls resistance to pathogen infection in Arabidopsis. Plant and Cell Physiology 59, 190–204.

12. Carbonell-Bejerano P., Diago M.P., Martínez-Abaigar J., Martínez-Zapater J.M., Tardáguila J. & Núñez-Olivera E. (2014) Solar ultraviolet radiation is necessary to enhance grapevine fruit ripening transcriptional and phenolic responses. BMC Plant Biology 14, 183.

13. Carra A., Mica E., Gambino G., Pindo M., Moser C., Pè M.E. & Schubert A. (2009) Cloning and characterization of small non-coding RNAs from grape. The Plant Journal 59, 750–763. doi:10.1111/j.1365-313X.2009.03906.x

14. Casati P. (2013) Analysis of UV-B regulated miRNAs and their targets in maize leaves. Plant Signaling & Behavior 10, e26758

15. Cavallini E., Matus J.T., Finezzo L., Zenoni S., Loyola R., Guzzo F.,…,Tornielli G.B. (2015) The phenylpropanoid pathway is controlled at different branches by a set of R2R3-MYB C2 repressors in grapevine. Plant Physiology 167, 1448–1470.

16. Cloix C., Kaiserli E., Heilmann M., Baxter K.J., Brown B.A., O’Hara A.,…,Jenkins GI. (2012) C-terminal region of the UV-B photoreceptor UVR8 initiates signaling through interaction with the COP1 protein. Proceedings of the National Academy of Sciences U.S.A. 109, 16366–16370.

17. Conn S., Curtin C., Bézier A., Franco C. & Zhang W. (2008) Purification, molecular cloning, and characterization of glutathione S-transferases (GSTs) from pigmented Vitis vinifera L. cell suspension cultures as putative anthocyanin transport proteins. Journal of Experimental Botany 59, 3621–3634.

18. Coruh C., Shahid S. & Axtell MJ. (2014) Seeing the forest for the trees: Annotating small RNA producing genes in plants. Current Opinion in Plant Biology 18, 87–95. doi: 10.1016/j.pbi.2014.02.008

19. Czemmel S., Höll J., Loyola R., Arce-Johnson P., Alcalde J.A., Matus J.T. & Bogs J. (2017) Transcriptome-wide identification of novel UV-B-and light modulated flavonol pathway genes controlled by VviMYBF1. Frontiers in Plant Science 8, 1084. doi.org/10.3389/fpls.2017.01084

20. Dauelsberg P., Matus J.T., Poupin M.J., Leiva-Ampuero A., Godoy F., Vega A. & Arce-Johnson P. (2011) Effect of pollination and fertilization on the expression of genes related to floral transition, hormone synthesis and berry development in grapevine. Journal of Plant Physiology 168, 1667–1674.

21. Ding D., Zhang L., Wang H., Liu Z., Zhang Z. & Zheng Y. (2009) Differential expression of miRNAs in response to salt stress in maize roots. Annals of Botany 103, 29–38.

22. Dotto M. & Casati P. (2017) Developmental reprogramming by UV-B radiation in plants. Plant Science 264, 96–101.

23. Elkayam E., Kuhn C.D., Tocilj A., Haase A.D., Greene E.M., Hannon G.J. & Joshua-Tor L. (2012) The structure of human Argonaute-2 in complex with miR-20a. Cell 150, 100–110.

24. Fahlgren N. & Carrington JC. (2010) miRNA target prediction in plants. In Meyers BC, Green PJ (Eds). Plant microRNAs. Methods in Molecular Biology 592, 51–57. Humana Press.

25. Fátyol K., Ludman M. & Burgyán J. (2016) Functional dissection of a plant Argonaute. Nucleic Acids Research 44, 1384–1397. doi:10.1093/nar/gkv1371.

26. Fei Q., Xia R. & Meyers B.C. (2013) Phased, secondary, small interfering RNAs in posttranscriptional regulatory networks. The Plant Cell 25, 2400–2415.

27. Frohnmeyer H., Loyall L., Blatt M.R. & Grabov A. (1999) Millisecond UV-B irradiation evokes prolonged elevation of cytosolic-free Ca2+ and stimulates gene expression in transgenic parsley cell cultures. The Plant Journal 20, 109–117.

28. Frohnmeyer H. & Staiger D. (2003) Ultraviolet-B radiation-mediated responses in plants. Balancing damage and protection. Plant Physiology 133, 1420–1428.

29. Gao F., Wang N., Li H., Liu J., Fu C., Xiao Z.,…,Zhou Y. (2016) Identification of drought-responsive microRNAs and their targets in Ammopiptanthus mongolicus by using high-throughput sequencing. Scientific Reports 6, 34601.

30. Geisler M., Frangne N., Gomes E., Martinoia E. & Palmgren MG. (2000) The ACA4 gene of Arabidopsis encodes a vacuolar membrane calcium pump that improves salt tolerance in yeast. Plant Physiology 124, 1814–1827.

31. Gill S.S., Anjum N.A., Gill R., Jha M. & Tuteja N. (2015) DNA damage and repair in plants under ultraviolet and ionizing radiations. Scientific World Journal 2015, 250158.

32. Gomez C., Conejero G., Torregrosa L., Cheynier V., Terrier N. & Ageorges A. (2011) In vivo grapevine anthocyanin transport involves vesicle-mediated trafficking and the contribution of anthoMATE transporters and GST. The Plant Journal 67, 960–970.

33. Gou J.Y., Felippes F.F., Liu C.J., Weigel D. & Wang J.W. (2011) Negative regulation of anthocyanin biosynthesis in Arabidopsis by a miR156-targeted SPL transcription factor. The Plant Cell 23, 1512–1522.

34. Gregory B.D., O’Malley R.C., Lister R., Urich M.A., Tonti-Filippini J., Chen H.,…,Ecker J.R. (2008) A link between RNA metabolism and silencing affecting Arabidopsis development. Developmental Cell 14, 854–866.

35. Guan X., Pang M., Nah G., Shi X., Ye W., Stelly D.M. & Chen Z.J. (2014) miR828 and miR858 regulate homoeologous gene functions in Arabidopsis trichome and cotton fibre development. Nature Communications 5, 3050.

36. Guo Q., Qu X. & Jin W. (2015) PhaseTank: genome-wide computational identification of phasiRNAs and their regulatory cascades. Bioinformatics 31, 284–286.

37. Gyula P., Baksa I., Tóth T., Irina Mohorianu I., Tamás Dalmay T. & Szittya G. (2018) Ambient temperature regulates the expression of a small set of sRNAs influencing plant development through NF-YA2 and YUC2. Plant, Cell and Environment online in advance, doi: 10.1111/pce.13355

38. Han L., Weng K., Ma H., Xiang G., Li Z., Wang Y.,…,Xu Y. (2016) Identification and characterization of Erysiphe necator-responsive microRNAs in Chinese wild Vitis pseudoreticulata by high-throughput sequencing. Frontiers in Plant Science 7, 621. doi: 10.3389/fpls.2016.00621

39. Harvey J.J., Lewsey M.G., Patel K., Westwood J., Heimstadt S., Carr J.P. & Baulcombe D.C. (2011) An antiviral defense role of AGO2 in plants. PLoS ONE 6, e14639.

40. Heijde M. & Ulm R. (2012) UV-B photoreceptor-mediated signalling in plants. Trends in Plant Science 17, 230–237.

41. Hiratsuka S., Onodera H., Kawai Y., Kubo T., Itoh H. & Wada R. (2001) ABA and sugar effects on anthocyanin formation in grape berry cultured in vitro. Scientia Horticulturae 90, 121–130.

42. Howell M.D., Fahlgren N., Chapman E.J., Cumbie J.S., Sullivan C.M., Givan S.A.,…,Carrington J.C. (2007) Genome-wide analysis of the RNA-DEPENDENT RNA POLYMERASE6/DICER-LIKE4 pathway in Arabidopsis reveals dependency on miRNA-and tasiRNA-directed targeting. The Plant Cell 19, 926–942.

43. Hsieh L.C., Lin S.I., Shih A.C., Chen J.W., Lin W.Y., Tseng C.Y.,…,Chiou T.J. (2009) Uncovering small RNA-mediated responses to phosphate deficiency in Arabidopsis by deep sequencing. Plant Physiology 151, 2120–2132.

44. Huang X., Ouyang X., Yang P., Lau O.S., Chen L., Wei N. & Deng X.W. (2013) Conversion from CUL4-based COP1-SPA E3 apparatus to UVR8-COP1-SPA complexes underlies a distinct biochemical function of COP1 under UV-B. Proceedings of the National Academy of Sciences U.S.A. 110, 16669–16674.

45. Jaillon O., Aury J.M., Noel B., Policriti A., Clepet C., Casagrande A.,…,Wincker P. (2007) The grapevine genome sequence suggests ancestral hexaploidization in major angiosperm phyla. Nature 449, 463–467.

46. Jenkins G.I. (2017) Photomorphogenic responses to ultraviolet-B light. Plant, Cell and Environment 40, 2544–2557.

47. Jia X., Ren L., Chen Q.J., Li R. & Tang G. (2009) UV-B-responsive microRNAs in Populus tremula. Journal of Plant Physiology 166, 2046–2057.

48. Johnson N.R., Yeoh J.M., Coruh C. & Axtell M.J. (2016) Improved placement of multi-mapping small RNAs. G3: Genes|Genomes|Genetics 6, 2103–2111.

49. Jung J.H., Seo Y.H., Seo P.J., Reyes J.L., Yun J., Chua N.H. & Park C.M. (2007) The GIGANTEA-regulated microRNA172 mediates photoperiodic flowering independent of CONSTANS in Arabidopsis. The Plant Cell 19, 2736–2748.

50. Kalvari I., Argasinska J., Quinones-Olvera N., Nawrocki E.P., Rivas E., Eddy S.R.,…,Petrov A.I. (2017) Rfam 13.0: shifting to a genome-centric resource for non-coding RNA families. Nucleic Acids Research 46, D335–D342.

51. Kawashima C.G., Matthewman C.A., Huang S., Lee B.R., Yoshimoto N., Koprivova A.,…,Kopriva S. (2011) Interplay of SLIM1 and miR395 in the regulation of sulfate assimilation in Arabidopsis. The Plant Journal 66, 863–876.

52. Koyama K., Sadamatsu K. & Goto-Yamamoto N. (2010) Abscisic acid stimulated ripening and gene expression in berry skins of the Cabernet Sauvignon grape. Functional and Integrative Genomics 10, 367–381.

53. Kozomara A. & Griffiths-Jones S. (2014) miRBase: annotating high confidence microRNAs using deep sequencing data. Nucleic Acids Research 42, D68–73.

54. Langmead B., Trapnell C., Pop M. & Salzberg S.L. (2009) Ultrafast and memory-efficient alignment of short DNA sequences to the human genome. Genome Biology 10, R25.

55. Lee E.K., Cibrian-Jaramillo A., Kolokotronis S-O., Katari M.S., Stamatakis A., Ott M.,…,DeSalle R. (2011) A functional phylogenomic view of the seed plants. PLoS Genetics 7, e1002411.

56. Lee J., He K., Stolc V., Lee H., Figueroa P., Gao Y., Tongprasit W.,…,Deng X.W. (2007) Analysis of transcription factor HY5 genomic binding sites revealed its hierarchical role in light regulation of development. The Plant Cell 19, 731–749.

57. Li F., Orban R. & Baker B. (2012) SoMART: a web server for plant miRNA, tasiRNA and target gene analysis. The Plant Journal 70, 891–901. doi: 10.1111/j.1365-313X.2012.04922.x

58. Li J., Ou-Lee T.M., Raba R., Amundson R.G. & Last R.L. (1993) Arabidopsis flavonoid mutants are hypersensitive to UV-B irradiation. The Plant Cell 5, 171–179.

59. Li S., Liu L., Zhuang X., Yu Y., Liu X., Cui X.,…,Chen X. (2013) microRNAs inhibit the translation of target mRNAs on the endoplasmic reticulum in Arabidopsis. Cell 153, 562–574.

60. Liang G., Li Y., He H., Wang F. & Yu D. (2013) Identification of miRNAs and miRNA-mediated regulatory pathways in Carica papaya. Planta 238, 739–752.

61. Lin S.I., Chiang S.F., Lin W.Y., Chen J.W., Tseng C.Y., Wu P.C. & Chiou T.J. (2008) Regulatory network of microRNA399 and PHO2 by systemic signaling. Plant Physiology 147, 732–746.

62. Liu F., Wang W., Sun X., Liang Z. & Wang F. (2015) Conserved and novel heat stress-responsive microRNAs were identified by deep sequencing in Saccharina japonica (Laminariales, Phaeophyta). Plant, Cell and Environment 38, 1357–1367.

63. Love M.I., Huber W. & Anders S. (2014) Moderated estimation of fold change and dispersion for RNA-seq data with DESeq2. Genome Biology 15, 550.

64. Loveys B.R. (1984) Abscisic acid transport and metabolism in grapevine (Vitis vinifera L.). New Phytologist 98, 575–582.

65. Loyola R., Herrera D., Mas A., Wong D.C., Höll J., Cavallini E.,…,Arce-Johnson P.. (2016) The photomorphogenic factors UV-B RECEPTOR 1, ELONGATED HYPOCOTYL 5, and HY5 HOMOLOGUE are part of the UV-B signalling pathway in grapevine and mediate flavonol accumulation in response to the environment. Journal of Experimental Botany 67, 5429–5445.

66. Lunde C., Zygadlo A., Simonsen H.T., Nielsen P.L., Blennow A. & Haldrup A. (2008) Sulfur starvation in rice: the effect on photosynthesis, carbohydrate metabolism, and oxidative stress protective pathways. Physiologia Plantarum 134, 508–521.

67. Luo Q-J., Mittal A., Jia F. & Rock C.D. (2012) An autoregulatory feedback loop involving PAP1 and TAS4 in response to sugars in Arabidopsis. Plant Molecular Biology 80, 117–129.

68. Luo Q-J., Samanta M.P., Köksal F., Janda J., Galbraith D.W., Richardson C.R.,…,Rock C.D. (2009) Evidence for antisense transcription associated with microRNA target mRNAs in Arabidopsis. PLoS Genetics 5, e1000457. doi 10.1371/journal.pgen.1000457

69. Mackerness S.A.H. (2000) Plant responses to UV-B (UV-B: 280–320 nm) stress: What are the key regulators? Plant Growth Regulation 37, 27–39

70. Martínez-Lüscher J., Morales F., Delrot S., Sanchez-Diaz M., Gomes E., Aguirreolea J. & Pascual I. (2013) Short-and long-term physiological responses of grapevine leaves to UV-B radiation. Plant Science 213, 114–122.

71. Massonnet M., Fasoli M., Battista Tornielli G., Altieri M., Sandri M., Zuccolotto P.,…,Pezzotti M. (2017) Ripening transcriptomic program in red and white grapevine varieties correlates with berry skin anthocyanin accumulation. Plant Physiology 174, 2376–2396.

72. Matus J.T., Cavallini E., Loyola R., Höll J., Finezzo L., Dal Santo S.,…,Arce-Johnson P. (2017) A group of grapevine MYBA transcription factors located in chromosome 14 control anthocyanin synthesis in vegetative organs with different specificities compared with the berry color locus. The Plant Journal 91, 220–236.

73. Matus J.T. (2016) Transcriptomic and metabolomic networks in the grape berry illustrate that it takes more than flavonoids to fight against ultraviolet radiation. Frontiers in Plant Science 7, 1337. doi: 10.3389/fpls.2016.01337

74. Mica E., Piccolo V., Delledonne M., Ferrarini A., Pezzotti M., Casati C.,…,Horner D.S. (2010) High throughput approaches reveal splicing of primary microRNA transcripts and tissue specific expression of mature microRNAs in Vitis vinifera. BMC Genomics 11, 109.

75. Montgomery T.A., Howell M.D., Cuperus J.T., Li D., Hansen J.E., Alexander A.L.,…Carrington J.C. (2008) Specificity of ARGONAUTE7-miR390 interaction and dual functionality in TAS3 trans-acting siRNA formation. Cell 133, 128–141.

76. Pagliarani C., Vitali M., Ferrero M., Vitulo N., Incarbone M., Lovisolo C.,…,Schubert A. (2017) The accumulation of miRNAs differentially modulated by drought stress is affected by grafting in grapevine. Plant Physiology 173, 2180–2195. doi: 10.1104/pp.16.01119

77. Papp I., Mur L.A., Dalmadi A., Dulai S. & Koncz C. (2004) A mutation in the Cap Binding Protein 20 gene confers drought tolerance to Arabidopsis. Plant Molecular Biology 55, 679–686

78. Pantaleo V., Szittya G., Moxon S., Miozzi L., Moulton V., Dalmay T. & Burgyan J. (2010) Identification of grapevine microRNAs and their targets using high-throughput sequencing and degradome analysis. The Plant Journal 62, 960–976.

79. Pantaleo V., Vitali M., Boccacci P., Miozzi L., Cuozzo D., Chitarra W.,…,Gambino G. (2016) Novel functional microRNAs from virus-free and infected Vitis vinifera plants under water stress. Scientific Reports 6, e20167. doi: 10.1038/srep20167

80. Paim-Pinto D.L., Brancadoro L., Dal Santo S., De Lorenzis G., Pezzotti M., Meyers B.C.,…,Mica E. (2016) The influence of genotype and environment on small RNA profiles in grapevine berry. Frontiers in Plant Science 7, 1459.

81. Pérez-Díaz R., Madrid-Espinoza J., Salinas-Cornejo J., González-Villanueva E. & Ruiz-Lara S. (2016) Differential roles for VviGST1, VviGST3, and VviGST4 in proanthocyanidin and anthocyanin transport in Vitis vinifera. Frontiers in Plant Science 7, 1166.

82. Pilati S., Bagagli G., Sonego P., Moretto M., Brazzale D., Castorina G.,…,Moser C. (2017) Abscisic acid is a major regulator of grape berry ripening onset: new insights into ABA signaling network. Frontiers in Plant Science 8, 1093. doi.org/10.3389/fpls.2017.01093

83. Piofczyk T., Jeena G. & Pecinka A. (2015) Arabidopsis thaliana natural variation reveals connections between UV radiation stress and plant pathogen-like defense responses. Plant Physiology and Biochemistry 93, 34–43.

84. Pitchiaya S., Heinicke L.A., Park J.I., Cameron E.L. & Walter N.G. (2017) Resolving subcellular miRNA trafficking and turnover at single-molecule resolution. Cell Reports 19, 630–642.

85. Piya S., Kihm C., Rice J.H., Baum T.J. & Hewezi T. (2017) Cooperative regulatory functions of miR858 and MYB83 during cyst nematode parasitism. Plant Physiology 174, 1897–1912.

86. Pontin M.A., Piccoli P.N., Francisco R., Bottini R., Martinez-Zapater J.M. & Lijavetzky D. (2010) Transcriptome changes in grapevine (Vitis vinifera L.) cv. Malbec leaves induced by ultraviolet-B radiation. BMC Plant Biology 10, 224

87. Rajagopalan R., Vaucheret H., Trejo J. & Bartel D.P. (2006) A diverse and evolutionarily fluid set of microRNAs in Arabidopsis thaliana. Genes & Development 20, 3407–3425.

88. Rajeswaran R. & Pooggin M.M. (2012) RDR6-mediated synthesis of complementary RNA is terminated by miRNA stably bound to template RNA. Nucleic Acids Research 40, 594–599.

89. Reid K.E., Olsson N., Schlosser J., Peng F. & Lund S.T. (2006) An optimized grapevine RNA isolation procedure and statistical determination of reference genes for real-time RTPCR during berry development. BMC Plant Biology 6, 27.

90. Ries G., Heller W., Puchta H., Sandermann H., Seidlitz H.K. & Hohn B. (2000) Elevated UV-B radiation reduces genome stability in plants. Nature 406, 98–101.

91. Rinaldo A.R., Cavallini E., Jia Y., Moss S.M., McDavid D.A., Hooper L.C.,…,Walker A.R. (2015) A grapevine anthocyanin acyltransferase, transcriptionally regulated by VvMYBA, can produce most acylated anthocyanins present in grape skins. Plant Physiology 169, 1897–1916.

92. Rizzini L., Favory J.J., Cloix C., Faggionato D., O’Hara A., Kaiserli E.,…,Ulm R. (2011) Perception of UV-B by the Arabidopsis UVR8 protein. Science 332, 103–106.

93. Rock C.D. (2013) Trans-acting small interfering RNA4: key to nutraceutical synthesis in grape development? Trends in Plant Science 18, 601–610.

94. Saijo Y., Sullivan J.A., Wang H., Yang J., Shen Y., Rubio V.,…,Deng X.W. (2003) The COP1-SPA1 interaction defines a critical step in phytochrome A-mediated regulation of HY5 activity. Genes & Development 17, 2642–2647.

95. Shin D.H., Choi M., Kim K., Bang G., Cho M., Choi S.B.,…,Park Y.I. (2013) HY5 regulates anthocyanin biosynthesis by inducing the transcriptional activation of the MYB75/PAP1 transcription factor in Arabidopsis. FEBS Letters 587, 1543–1547.

96. Shivaprasad P.V., Chen H.M., Patel K., Bond D.M., Santos B.A. & Baulcombe D.C. (2012) A microRNA superfamily regulates nucleotide binding site-leucine-rich repeats and other mRNAs. The Plant Cell 24, 859–874.

97. Shao F. & Lu S. (2013) Genome-wide identification, molecular cloning, expression profiling and posttranscriptional regulation analysis of the Argonaute gene family in Salvia miltiorrhiza, an emerging model medicinal plant. BMC Genomics 14, 512.

98. Si-Ammour A., Windels D., Arn-Bouldoires E., Kutter C., Ailhas J., Meins F. & Vazquez F. (2011) miR393 and secondary siRNAs regulate expression of the TIR1/AFB2 auxin receptor clade and auxin-related development of Arabidopsis leaves. Plant Physiology 157, 683–691.

99. Solofoharivelo M.C., van der Walt A.P., Stephan D., Burger J.T. & Murray S.L. (2014) MicroRNAs in fruit trees: discovery, diversity and future research directions. Plant Biology 16, 856–865.

100. Speth C., Willing E.M., Rausch S., Schneeberger K. & Laubinger S. (2013) RACK1 scaffold proteins influence miRNA abundance in Arabidopsis. The Plant Journal 76, 433–445.

101. Stracke R., Favory J.J., Gruber H., Bartelniewoehner L., Bartels S., Binkert M.,…,Ulm R. (2010) The Arabidopsis bZIP transcription factor HY5 regulates expression of the PFG1/MYB12 gene in response to light and ultraviolet-B radiation. Plant, Cell and Environment 33, 88–103.

102. Sun X., Fan G., Su L., Wang W., Liang Z., Li S. & Xin H. (2015) Identification of cold-inducible microRNAs in grapevine. Frontiers in Plant Science 6, 595.

103. Suzuki M., Nakabayashi R., Ogata Y., Sakurai N., Tokimatsu T., Goto S.,…,Shiratake K. (2015) Multiomics in grape berry skin revealed specific induction of the stilbene synthetic pathway by ultraviolet-C irradiation. Plant Physiology 168, 47–59. doi:10.1104/pp.114.254375.

104. Terrier N., Glissant D., Grimplet J., Barrieu F., Abbal P., Couture C.,…,Hamdi S. (2005) Isogene specific oligo arrays reveal multifaceted changes in gene expression during grape berry (Vitis vinifera L.) development. Planta 222, 832–847. 105.

105. Tilbrook K., Dubois M., Crocco C.D., Yin R., Chappuis R., Allorent G.,…,Ulm R. (2016) UV-B perception and acclimation in Chlamydomonas reinhardtii. The Plant Cell 28, 966–983.

106. Tuteja N., Singh M.B., Misra M.K., Bhalla P.L. & Tuteja R. (2001) Molecular mechanisms of DNA damage and repair: progress in plants. Critical Reviews in Biochemistry and Molecular Biology 36, 337–397.

107. Valoczi A., Hornyik C., Varga N., Burgyan J., Kauppinen S. & Havelda Z. (2004) Sensitive and specific detection of microRNAs by northern blot analysis using LNA-modified oligonucleotide probes. Nucleic Acids Research 32, e175.

108. Vanhaelewyn L., Prinsen E., Van Der Straeten D. & Vandenbussche F. (2016) Hormone-controlled UV-B responses in plants. Journal of Experimental Botany 67, 4469–4482.

109. Wagoner J.A., Sun T., Lin L. & Hanson M.R. (2014) Cytidine deaminase motifs within the DYW domain of two pentatricopeptide repeat-containing proteins are required for site-specific chloroplast RNA editing. Journal of Biological Chemistry 290, 2957–2968. doi: 10.1074/jbc.M114.622084

110. Walker A.R., Lee E., Bogs J., McDavid D.A., Thomas M.R. & Robinson S.P. (2007) White grapes arose through the mutation of two similar and adjacent regulatory genes. The Plant Journal 49, 772–785.

111. Wang B., Sun Y.F., Song N., Wang X.J., Feng H., Huang L.L. & Kang Z.S. (2013) Identification of UV-B-induced microRNAs in wheat. Genetics and Molecular Research 12, 4213–4221.

112. Wang C., Han J., Liu C., Kibet K.N., Kayesh E., Shangguan L.,…,Fang J. (2012) Identification of microRNAs from Amur grape (Vitis amurensis Rupr.) by deep sequencing and analysis of microRNA variations with bioinformatics. BMC Genomics 13, 122.

113. Wang C., Leng X., Zhang Y., Kayesh E., Zhang Y., Sun X. & Fang J. (2014) Transcriptome-wide analysis of dynamic variations in regulation modes of grapevine microRNAs on their target genes during grapevine development. Plant Molecular Biology 84, 269–285.

114. Wang Y., Itaya A., Zhong X., Wu Y., Zhang J., van der Knaap E.,…,Ding B. (2011) Function and evolution of a microRNA that regulates a Ca2+-ATPase and triggers the formation of phased small interfering RNAs in tomato reproductive growth. The Plant Cell 23, 3185–3203.

115. Wei W., Ba Z., Gao M., Wu Y., Ma Y., Amiard S.,…,Qi Y. (2012) A role for small RNAs in DNA double-strand break repair. Cell 149, 101–112.

116. Wu D., Hu Q., Yan Z., Chen W., Yan C., Huang X.,…,Shi Y. (2012) Structural basis of ultraviolet-B perception by UVR8. Nature 484, 214–219.

117. Xia R., Zhu H., An Y-q., Beers E.J. & Liu Z. 2012. Apple miRNAs and tasiRNAs with novel regulatory networks. Genome Biology 13, R47.

118. Xia R., Meyers B.C., Liu Z., Beers E.P. & Ye S. (2013) MicroRNA superfamilies descended from miR390 and their roles in secondary small interfering RNA biogenesis in eudicots. The Plant Cell 25, 1555–1572.

119. Yamakawa T., Kato S., Ishida K., Kodama T. & Minoda Y. (1983) Production of anthocyanins by Vitis cells is suspension culture. Agricultural and Biological Chemistry 47, 2185–2191.

120. Yi X., Zhang Z., Ling Y., Xu W. & Su Z. (2015) PNRD: a plant non-coding RNA database. Nucleic Acids Research 43, D982–989.

121. Yoon M.Y., Kim M.Y., Shim S., Kim K.D., Ha J., Shin J.H.,…,Lee S.H. (2016) Transcriptomic profiling of soybean in response to high-intensity UV-B irradiation reveals stress defense signaling. Frontiers in Plant Science 7, 1917.

122. Zhai J., Jeong D.H., De Paoli E., Park S., Rosen B.D., Li Y.,…,Meyers B.C. (2011) MicroRNAs as master regulators of the plant NB-LRR defense gene family via the production of phased, trans-acting siRNAs. Genes & Development 25, 2540–2553. doi: 10.1101/gad.177527.111

123. Zhang C., Li G., Wang J. & Fang J. (2012) Identification of trans-acting siRNAs and their regulatory cascades in grapevine. Bioinformatics 28, 2561–2568.

124. Zhang H., Zhao X., Li J., Cai H., Deng X.W. & Li L. (2014) MicroRNA408 is critical for the HY5-SPL7 gene network that mediates the coordinated response to light and copper. The Plant Cell 26, 4933–4953.

125. Zhang X., Niu D.D., Carbonell A., Wang A., Lee A., Tun V.,…,Jin H. (2014) ARGONAUTE PIWI domain and microRNA duplex structure regulate small RNA sorting in Arabidopsis. Nature Communications 5, 5468.

126. Zhang Z.L., Ogawa M., Fleet C.M., Zentella R., Hu J., Heo J.O.,…,Sun T.P. (2011) Scarecrow-like 3 promotes gibberellin signaling by antagonizing master growth repressor DELLA in Arabidopsis. Proceedings of the National Academy of Sciences U.S.A. 108, 2160–2165.

127. Zhou X., Wang G. & Zhang W. (2007) UV-B responsive microRNA genes in Arabidopsis thaliana. Molecular Systems Biology 3, 103. doi: 10.1038/msb4100143

128. Zhou L., Liu Y., Liu Z., Kong D., Duan M. & Luo L. (2010) Genome-wide identification and analysis of drought-responsive microRNAs in Oryza sativa. Journal of Experimental Botany 61, 4157–4168.

129. Zhou B., Fan P., Li Y., Yan H. & Xu Q. (2016) Exploring miRNAs involved in blue/UV-A light response in Brassica rapa reveals special regulatory mode during seedling development. BMC Plant Biology 16, 111.

130. Zhu H., Xia R., Zhao B., An Y-q., Dardick C.D., Callahan A.M. & Liu Z. (2012) Unique expression, processing regulation, and regulatory network of peach (Prunus persica) miRNAs. BMC Plant Biology 12, 149.

